# Predicting gene regulatory networks from cell atlases

**DOI:** 10.1101/2020.08.21.261735

**Authors:** Andreas Fønss Møller, Kedar Nath Natarajan

**Affiliations:** Functional Genomics and Metabolism Unit, Department of Biochemistry and Molecular Biology, University of Southern Denmark, Denmark; Danish Institute of Advanced Study (D-IAS), University of Southern Denmark, Denmark

**Keywords:** Single-cell RNA-sequencing, Mouse cell atlas, Regulons, Gene regulatory networks

## Abstract

Recent single-cell RNA-sequencing atlases have surveyed and identified major cell-types across different mouse tissues. Here, we computationally reconstruct gene regulatory networks from 3 major mouse cell atlases to capture functional regulators critical for cell identity, while accounting for a variety of technical differences including sampled tissues, sequencing depth and author assigned cell-type labels. Extracting the regulatory crosstalk from mouse atlases, we identify and distinguish global regulons active in multiple cell-types from specialised cell-type specific regulons. We demonstrate that regulon activities accurately distinguish individual cell types, despite differences between individual atlases. We generate an integrated network that further uncovers regulon modules with coordinated activities critical for cell-types, and validate modules using available experimental data. Inferring regulatory networks during myeloid differentiation from wildtype and Irf8 KO cells, we uncover functional contribution of Irf8 regulon activity and composition towards monocyte lineage. Our analysis provides an avenue to further extract and integrate the regulatory crosstalk from single-cell expression data.

**Summary:** Integrated single-cell gene regulatory network from three mouse cell atlases captures global and cell-type specific regulatory modules and crosstalk, important for cellular identity.

## Introduction

Multi-cellular organisms are composed of different tissues consisting of varied cell-types that are regulated at single-cell level. Single-cell RNA-sequencing (scRNA-seq) enables high-throughput gene expression measurements for unbiased and comprehensive classification of cell-types and factors that contribute to individual cell states [1,2]. The underlying expression heterogeneity between single-cells can be attributed to finer grouping of cell-types, inherent stochasticity and variations in underlying functional and regulatory crosstalk [3–6]. Single-cells maintain their cell state and also respond to a variety of external cues by modulating transcriptional changes, which are governed by complex gene-regulatory networks (GRNs) [7,8]. A GRN is specific combination of transcription factors (TFs) and co-factors that interact with cis-regulatory genomic regions to mediate a specialised transcriptional programme within individual cells [9,10]. Briefly, a regulon is collection of a transcription factor (TF) and all its transcriptional target genes. The GRNs define and govern individual cell-type definition, transcriptional states, spatial patterning and responses to signalling, cell fate cues [11]. Recent computational approaches have enabled inference of the gene regulatory circuitry from scRNA-seq datasets [9,12–16].

Recently two major single-cell mouse atlases studies were published [17,18]. The Tabula Muris (TM) and Mouse Cell Atlas (MCA), profiled >500k individual single-cells using three different scRNA-seq platforms, across multiple murine tissues to provide a broad survey of constituent cell-types and gene expression patterns and thereby demarcating shared and unique signatures across single-cells. The 3 cell atlases utilize different scRNA-seq platforms and technologies including Smart-seq2 (TM-SS2: [19], 10x Chromium (TM-10x: [20] and Microwell-seq [18].

For regulatory and mechanistic insights beyond cell-type survey across the 3 atlases, we have to extend analysis beyond comparison of gene expression patterns. The computational inference of TFs and their regulated gene sets (‘regulons’) provides an avenue to extract the regulatory crosstalk from single-cell expression data [9,10,21,22]. Here, we set out to comprehensively reconstruct GRNs from single-cell mouse cell atlases and address the following questions: (i) Which TFs, master regulators and co-factors (i.e. regulons) govern tissue and cell-types (ii) Do inferred regulons regulate ‘specific’ or multiple cell-types. (iii) which regulons and regulated gene sets are critical for individual cell identity?

In our integrative analysis, we identify regulon modules that globally regulate multiple cell groups and tissues across cell atlases. The cell-type specific regulons are characterised by distinct composition and activity, critical for their definition. We find that regulons and their activity scores are robust indicators of cell-type identity across cell atlases, irrespective of composition differences. We uncover modules of regulons and reconstruct an integrated atlas-scale regulatory network, and also validate network interactions using available experimental datasets. Importantly, we uncover the functional consequence of Irf8 regulon perturbation at single-cell level during myeloid lineage decisions from wildtype and Irf8 knockout cells. We uncover a distinctly depleted Irf8 regulon composition and activity Irf8 knockouts, validating the specification bias from monocytes to granulocytes. This work provides a consensus view of key regulators functioning in different cell-types that define cellular programs at single-cell level.

## Results

To identify regulatory networks across the different mouse cell-types and tissues, we analysed both ‘Tabula Muris’ and ‘Mouse Cell Atlas’ (MCA) scRNA-seq studies [17,18]. The Tabula Muris contains >130k annotated single-cells profiled using two scRNA-seq methods (referred as atlases), full-length Smart-seq2 (~54k single cells, 18 tissues, 81 cell-types) and 3’-end droplet based 10x Chromium (70k single cells, 12 tissues, 55 cell-types). The MCA contains >230k annotated single cells profiled using the authors 3’-end microwell-seq platform (38 tissues, 760 cell-types; Supplementary methods).

We aimed to integrate the atlases to identify cell-type specific regulons and build a consensus regulon atlas (Fig 1A; Detailed workflow in Fig. S1). As each atlas samples different mouse tissues and scRNA-seq technologies (full length vs 3’end) to identify hundreds of varied cell types across cellular resolutions (discussed below), a fundamental challenge is to effectively link the original author’s cell-type annotation across cell atlases. We address the challenge of integrating cell-type classification by combining two complementary approaches. Firstly, we manually devised a generalised vocabulary consisting of broadly defined ‘7 cell groups’ for an standardise annotation between cell atlases (3 datasets). Secondly, we utilise scMAP, an unsupervised scRNA-seq cell projection method [23], to link the original author’s cell-type annotation across cell atlases (Supplementary methods). By utilizing Tabula Muris 10x (TM-10x) Chromium annotations as a reference and by combining both approaches, our generalised vocabulary contains ‘7 cell groups’ consisting of ‘55 reference cell-types’. The 7 cell groups include *Immune* (22 subgroups), *Specialised* (12 subgroups), *Epithelial* (7 subgroups), *Stem* (4 subgroups), *Endothelial* (4 subgroups), *Basal* (3 subgroups) and *Blood* (3 subgroups) (Fig. S2A). Subsequently, we applied our two-step approach to individual atlases i.e. TM-10x (Fig. S2B), Tabula Muris Smart-seq2 (TM-SS2; Fig. S3A), MCA (Fig. S3A-B) and to all atlases integrated together (Fig. S4A). Our approach allows us to build and link an integrated mouse atlas consisting of 831-author assigned unique cell-type labels from 50 tissues to a consensus of 55 reference cell-types and 7 cell groups (Fig. S4A, methods and Table S1).

**Figure 1:**
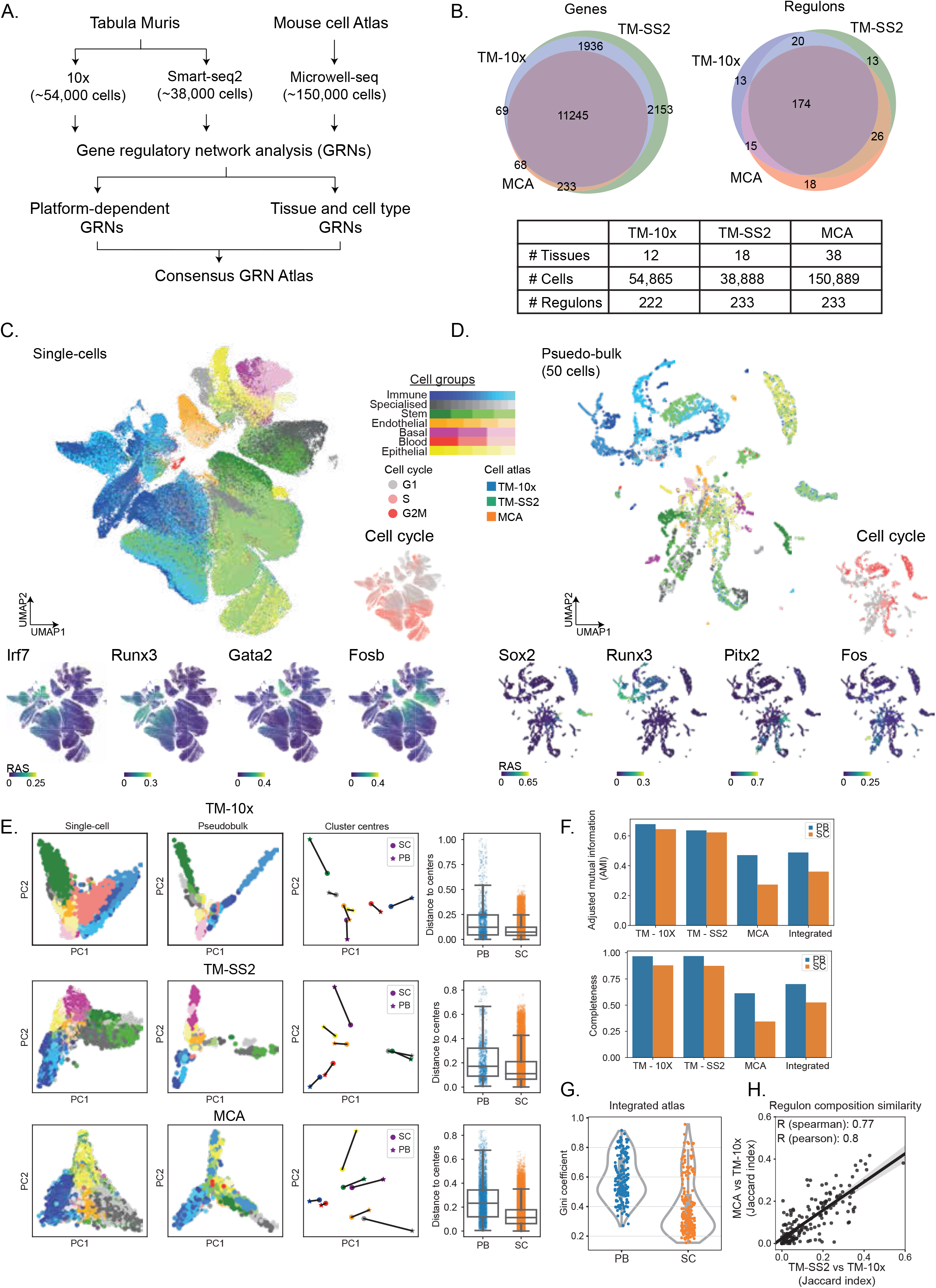
**A.** Overview of datasets and analysis performed in this study. **B.** Venn plots and table representation of shared and unique features across cell atlases including tissues, number of cells and regulons across cell. We used 11,245 overlapping genes and resulting 279 unique regulons for regulatory analysis. **C.** UMAP embedding of single-cells (centre) based on regulon activity scores (RAS) from integrated mouse atlases. The individual cells are coloured by 55 reference cell types corresponding to 7 cell groups. The surrounding plots highlight examples of individual regulons (Irf7, Runx3, Gata2, Fosb) coloured by RAS, predicted cell cycle stages (right), and overlaid on UMAP. **D.** UMAP embedding of 50-cell pseudobulk samples, based on regulon activity scores (RAS) from integrated mouse atlases. The surrounding plots highlight examples of individual regulons (Sox2, Runx3, Pitx2, Fos) coloured by RAS, predicted cell cycle stages (right), and overlaid on UMAP. The pseudobulk is generated by averaging expression of 50 cells across same tissues, using author assigned tissue and cell-type labels; and performing SCENIC regulatory inference. **E.** Principal component analysis (PCA) of matched single- and pseudobulk cells based on RAS across individual atlases and coloured by 7 cell groups (first two columns). For each of the 7 cell groups, we plotted cluster centroid (column 3) and connected single- (circles) and pseudobulk (asterisk). Box plots (column 4) represent Euclidean distance of individual single- and pseudobulk cells to respective cell group centroid. The distance is a measure of clustering i.e., pseudobulk cells are more separated with increased distance to cluster center than singlecells. **F.** Different measures of cluster comparison (top: Adjusted mutual information, bottom: Completeness) between pseudobulk and single-cells across integrated and individual mouse atlases, considering 7 cell groups **G.** Distribution of Gini coefficients per regulon in pseudobulk and single-cells across integrated atlas, considering all 7 cell groups. The Gini coefficient is a measure of inequality i.e. whether individual regulons contribute to individual (smaller Gini) or multiple cell groups (higher Gini). The pseudobulk cells have higher Gini coefficients and tighter distributions compared to single-cells, which highlights their contribution to effectively distinguish multiple cell groups. **H.** Comparison of regulon composition between atlases (Pairwise Jaccard index) considering TM-10x as reference. Each dot represents a regulon and overlap of its target genes across 3 atlases. The shaded area represents 95% confidence interval from the linear regression line.

**Figure 2:**
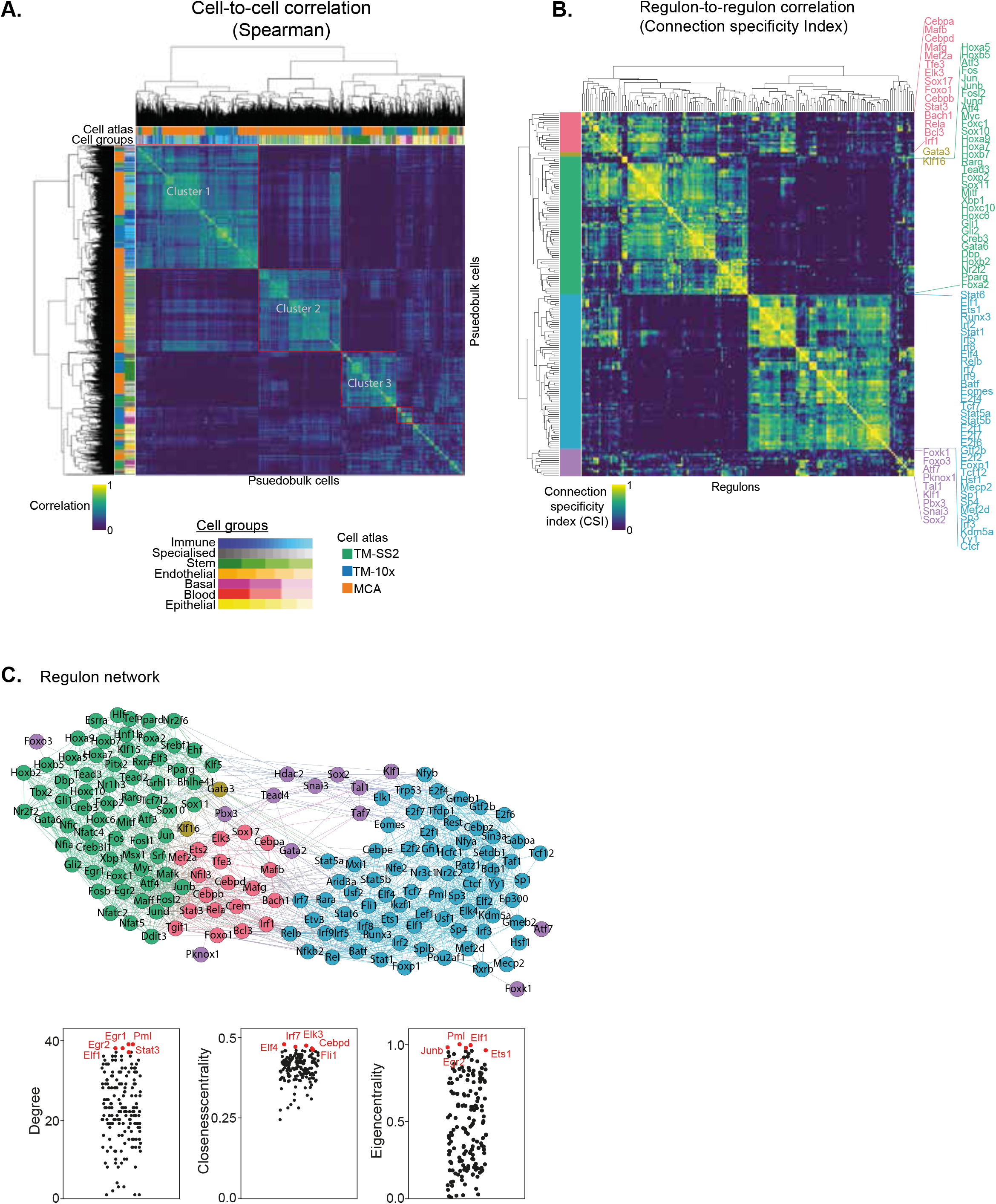
**A.** Spearman cell-to-cell correlation map across three atlases. The first column (and topmost row) indicates the respective mouse atlases, while second column (and second row) indicate the 55 reference cell types. The clusters are highlighted in red rectangles. **B.** Connection specificity index (CSI) matrix highlights regulon-to-regulon correlation in pseudo-bulk cells across integrated atlas. Hierarchical clustering of regulons identifies 5 distinct regulon modules (first column), which capture both global and distinct regulatory roles across cell groups and tissues. Selected regulons are coloured by module and listed next to heatmap. **C.** Undirected regulon network generated from strongly correlated CSI scores (Fig. 2B). Each regulon is represented as a node, and regulons pairs with strongest associated interactions (CSI scores > 0.7) are connected with undirected and unweighted edges. The larger modules 3 (green) and 4 (blue) are bridged by smaller modules. Bottom: Examples of individual regulons contributing to different network features (Degree, Closeness centrality and Eigen centrality).

We support the robustness of our generalised vocabulary and projection mapping approach by multiple analysis. Across individual tissues, we re-confirmed that author cell type labels are robustly mapped to reference cell types and cell groups both in individual and integrated atlas (Fig. S5A Liver, Fig. S5B Spleen, S2A, S3A-B). The individual atlases have technical difference owing to different number of cells profiled (S6A top panel), sequencing depth (library size, S6A middle panel), number of tissues profiled (12 TM-10x, 18 TM-SS2, 38 MCA; S2B, S3A-B), scRNA-seq chemistry (Full-length vs 3’), scRNA-seq platform and number of genes detected (S6A bottom panel). The dropout distribution for individual atlases highlights the relationship between number of cells profiled, library size and genes detected (Fig. S6B). Specifically, MCA compared to Tabula Muris atlases has the highest number of profiled cells at sparse sequencing depth, lower gene detected and highest dropout rates across reference cell groups (S6A-B). Our 7 reference cell groups have high and proportional number of cells from both integrated (Fig. S6C) and individual atlas (Fig. S6D). For example, the Immune cell group consists of 20,133 individual cells classified across 22 reference cell types, while the blood cell group consists of 1559 cells classified into 3 reference cell types (Fig. S6C and S2A). We further present the different technical features for each reference cell type across integrated and individual atlas (Fig. S7A). Our two-step approach consisting of simplified cell group and subgroup classification allows us to mitigate technical and cell-type label discrepancies, integrate mouse cell atlases to investigate global and specific regulators across atlases.

Feature selection is a crucial aspect for robust regulon inference and composition. We tested a variety of different feature sets for both integrated and individual cell atlases. We selected a reasonable cut-off of genes detected in at least 10% of all single-cells, consisting of 11,245 overlapping genes across three atlases (Fig 1B). This cutoff robustly and proportionally captures the reference cell groups across integrated and individual atlases, despite the technical differences (Fig. S7). To infer gene regulatory networks (GRNs), we applied SCENIC, a framework for network inference, reconstruction and clustering from scRNA-seq data [10]. The SCENIC framework is applied directly on single-cell expression matrix and combines three different approaches for regulon identification and activity. The three approaches consist of (i) ‘GRNBoost’ for identification of TFs and co-expressed genes from single-cell expression matrix, (ii) ‘RcisTarget’ for defining ‘regulons’ (i.e enriched and validated TFs with their direct downstream target genes containing annotated motif, and prunes co-expressed indirect targets), (iii) ‘AUCell’ for scoring regulon activity (RAS: regulon activity scores) in single-cells. Our motivation for using SCENIC for atlas scale regulon inference was three-fold. Firstly, SCENIC utilises GRNBoost, RCisTargets and AUCell to identify and score known, direct TF-target interactions while pruning away indirect and co-expressed TF-target links. The RCisTarget crossmatches regulons with known TF-target databases, as opposed to de-novo predictions. This allows inference of both TF-TF and TF-target relationships and scoring of TF-target relationships, unlike other approaches. Secondly, SCENIC does not require a single-cell trajectory/pseudotime, unlike other widely applied GRN methods [12], and therefore is well suited to cell atlas-scale analysis. Thirdly, SCENIC is widely used and GENIE3/GRNBoost are scored amongst the top reconstruction methods in recent benchmarking study [12]. Applying SCENIC, we identify 279 unique regulons, with >60% (174 regulons) shared across the three atlases (Fig 1B). The high degree of regulon overlap between the three atlases, in spite of technical differences highlights that single-cell regulatory state is predominantly governed by core set regulators and their activities within individual cells. A recent study also applied SCENIC, but only for MCA data using only the author assigned cell-type labels [21].

To distinguish the regulatory activity within individual cells, we performed dimensionality reduction using UMAP on regulon activity scores (RAS) of ~250k single-cells, integrating all atlases [24]. We coloured individual cells using the reference cell groups (Fig 1C), predicted cell cycle stage (Fig 1C, right) and tissue of origin (Fig. S8A). We observe good visual separation between the 7 cell groups based on RAS, highlighting robustness of cell group classification and ability of RAS to distinguish functional cell-types in integrated atlas. The overlapping cell groups are biologically and functionally related, with similar RAS and tissue origin. For example, subset of Immune (blue) and Stem (green) cell groups originating from bone marrow overlap in the integrated atlas (bottom left: Fig 1C, Fig S8A). The cell cycle stage prediction based on scRNA-seq is also consistent with cell groups and reference cell-type classification [25,26]. As expected, most Stem and Immune reference cell-types are actively cycling (S, G2M stage; Fig 1C), while subsets of Specialised, Stem cell-types are in G0/G1 stage originating from brain, liver and bone marrow. We could further classify Immune cell groups into proliferating (i.e. T-cells from spleen) and quiescent (grey G0/G1 monocytes). Furthermore, both endothelial cells and hepatocytes are in the G1 stage, while erythroblasts are actively cycling. We next focussed on both global regulons active across multiple cell groups and cell-type specific regulons within the integrated atlas. The Irf7 (2437 unique genes) and Runx3 (474 unique genes) are enriched in the Immune cell group (Fig 1C) [27,28]. The general transcription factor E2F4 is enriched across most proliferating cells, while E2F7 (an atypical E2F transcription factor) is exclusively active in a subset of highly proliferating cell groups (Fig. S8C). The Foxo1 and Cebpe regulons are also enriched across multiple cell groups (Immune, Stem and Epithelial). The specific and enriched regulons include Fosb (1352 unique genes; Endothelial and Stem), Gata2 regulon (1594 unique genes; Endothelial) and Gli1 (114 unique genes; Bladder cells within Specialised) (Fig 1C, S8C), Sox17 (267 unique genes; Endothelial) [29] and Cebpa (1201 unique genes; Pancreas and myeloid single-cells within Immune cell group; Fig S8C). The individual regulons and their compositions are detailed in Table S2.

The GRN inference on ~250k unevenly sampled single-cells is computationally intensive and also impacted by scRNA-seq platform-specific biases (Fig. S6A-B, S7). To address this, we generated pseudobulk cells by averaging scRNA-seq expression over 50 cells. The pseudobulk approach is computationally robust and also accounts for technical differences between atlases (additional comparison below). We re-performed the SCENIC framework on pseudobulk cells across the integrated atlas, projected individual cells on UMAP based on RAS, coloured by cell groups (Fig 1D), predicted cell cycle stage (Fig 1D) and tissue of origin (Fig. S8B). We expected a better separation with pseudobulk owing to reduced technical noise (50 cell average) and more robust RAS. Consistently, the cell-type separation is visually refined, with a strong overlap of cell groups across different tissues (Fig. S8B) and recovery of both general and specific regulons. These include Runx3 (Immune), Sox2 (752 genes; Stem and Immune) [30], Homeodomain Pitx2 (63 unique genes from bladder, skin and heart), Atf3, Fos (Basal) and Foxc1 (564 unique genes; Stem) (Fig 1D, S7B, D).

We next assessed regulon activities in individual cell atlases by re-performing SCENIC (regulon scoring by AUCell) and compared with integrated mouse atlas. The UMAP embedding based on RAS distinctly separates cell groups within individual atlases, in both single- (Fig. S9A top panel) and pseudobulk cells (S9B top panel). The MCA dataset has the largest number of cells, increased technical noise, lower gene detection (Fig. S6A bottom row), and is enriched for Immune and Stem cell groups. Consequently, the MCA single-cell UMAP partially distinguishes reference cell groups compared to other atlases (Fig. S9A top right). However, the MCA pseudobulk UMAP clearly resolves cell groups, while retaining robust regulon activities (Fig. S9B). Across individual atlases, we recapitulate several integrated atlas features including global and cell group specific regulons. For example, the large Irf8 (2988 unique genes) and smaller Tcf7 regulons (25 unique genes) are both highly specific and enriched in Immune across multiple tissues in all atlases (Fig. S9A-B, S10A-B). Within individual atlases, we also observe finer celltype and tissue specific regulon activity including Sox17 (267 genes), Sox2 (752 genes) and Pparg (584 genes) (Fig. S9A). We also observe better reference cell types mixing originating from similar tissues in pseudobulk compared to single-cells (Fig. S10A-B). The individual regulons and mean RAS for reference cell types is reported in Table S3.

To highlight that pseudobulk robustly captures regulon activities across cell groups in comparison to single-cell, we performed several quantitative and qualitative comparisons. First, we distinguish single- and pseudobulk cells by Principal component analysis (PCA) for each atlas, coloured by 7 cell groups (Fig. 1E, methods). The pseudobulk cells are better separated than single-cells by PCA, but not as distinctly as with non-linear methods (e.g. UMAP; Fig. 1CD). We next compare the distances of individual single- and pseudobulk cells to cluster centers of 7 reference cell groups. Globally, the pseudobulk cells have increased distance to cluster center than single-cells, indicating a more homogeneous separation and increased cell group resolution based on RAS (Fig. 1E). To compare the clustering of cell groups between pseudobulk and single-cells, we computed Adjusted Mutual information (AMI) and Completeness (Fig. 1F). The AMI score is a symmetric measure of the agreement between two independent clustering labels i.e. pseudobulk and single-cells given the reference cell group labels, while Completeness compares clustering given a ground truth by measuring the membership of data points to the same cluster. Across both individual and integrated atlas, the AMI scores are consistently higher in pseudobulk than single-cells (Fig. 1F top). Notably, the MCA AMI is significantly lower than other atlases, reflecting the poorer cell group separation in single-cells compared to pseudobulk (Fig. S9A-B top right). We calculate Completeness measure between pseudobulk and single-cells by comparing k-means clustering (k=7) to our reference 7 cell groups across both individual and integrated atlas (Fig. 1F bottom). To measure the importance of regulons in driving integrated atlas, we computed Gini coefficient for each regulon (using RAS) across pseudobulk and singlecells. The Gini coefficient is a measure of equality in a given distribution, i.e. whether individual regulons drive all cell groups (Gini=0; complete equality) or multiple regulons drive most cell groups (Gini=1; inequality). Across integrated atlas, the pseudobulk have a higher median Gini coefficient with narrow dispersion compared to single-cells (Fig. 1G). Notably, the single-cells RAS tend to be skewed towards lower Gini coefficient, consistent with poorer separation of cell groups in lower dimensions (UMAP and PCA; Fig. 1C-D), compared to pseudobulk. We observe the same trend of Gini coefficients across individual cell groups (Fig. S11A). To compare and validate the clustering between integrated and individual atlases across single- and pseudoubulk cells, we compute Silhouette score (Fig. S11B). The Silhouette score is a measure of similarity between different clustering and considers both cohesion (within clusters) and separation (distance between clusters). We observe a positive Silhouette score for both integrated and individual atlases, with higher scores in pseudobulk cells. Consistent with previous observations, the MCA pseudobulk has significantly improved clustering and Silhouette scores compared to single-cells. Additionally, we assess the regulon composition similarity between pseudoubulk and single-cells by pairwise atlas comparison and computing Jaccard similarity index (Fig. 1H). The Jaccard index is strongly correlated (R_pearson_=0.8), highlighting that target gene compositions are similar in individual atlases (Individual regulon examples described in Fig. 3A-D and Fig. S18-20). Lastly, we compare RAS between single- and pseudobulk cells and observed significantly improved correlation in individual cell groups (Fig. S11C)

**Figure 3:**
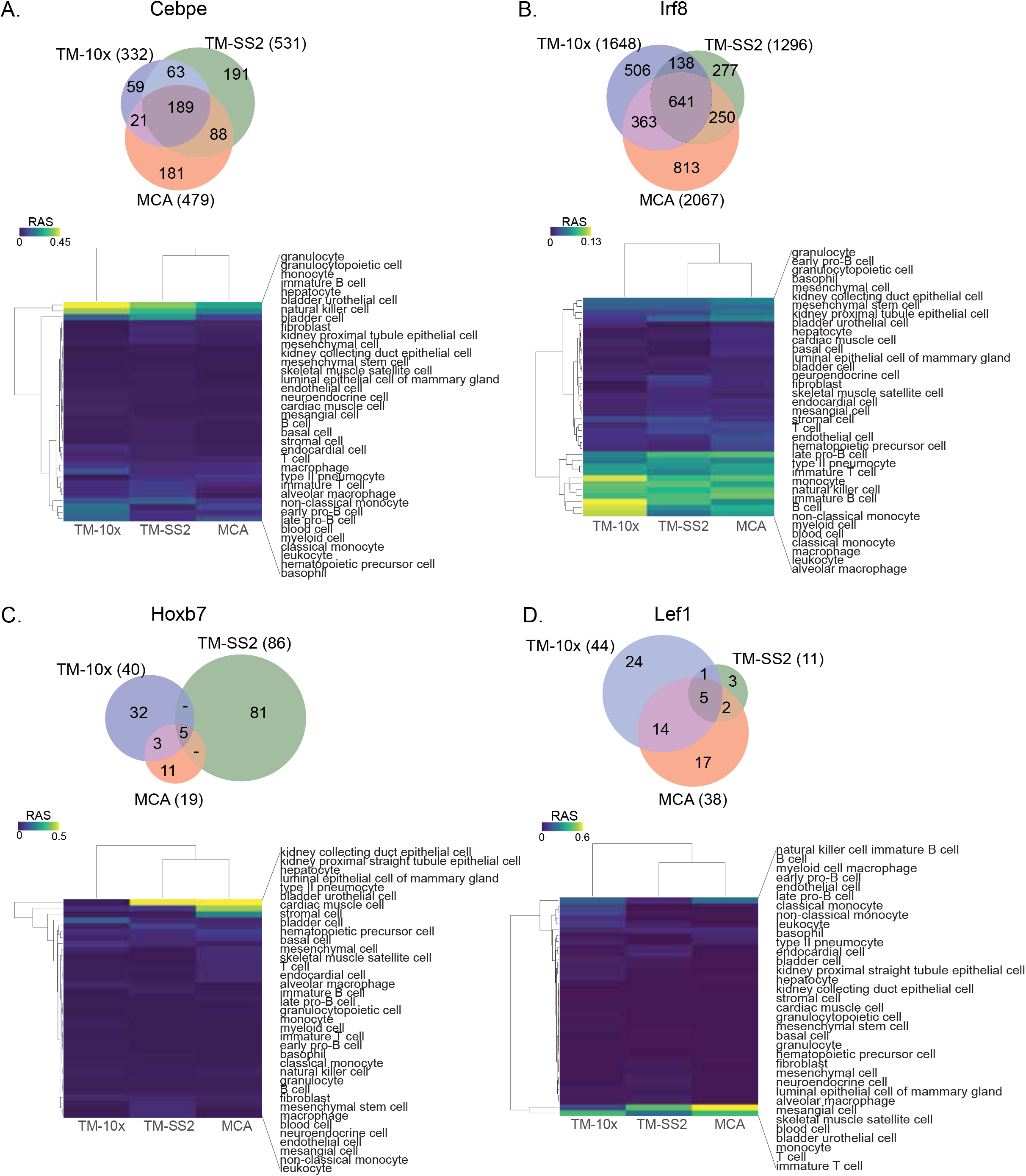
**A-D.** Venn plots of representative individual regulons, gene compositions, overlap across individual atlas and specific cell type regulation (**A**) Irf8, (**B**) Irf8, (**C**) Hoxb7 and (**B**) Lef1. The heatmap represent z-scaled mean RAS across different cell types.

Given the different technical differences between individual atlases (Dropouts, tissues profiled, scRNA-seq protocol, sequencing depth etc.,), we also assessed whether batch effects confound RAS across mouse atlases. Although SCENIC analysis has been shown to be unaffected by batch and technical effects [10], we performed batch correction on a common tissue (Spleen) profiled by both TM-10x and TM-SS2 atlases. We apply two methods ‘Batch-balanced KNN’ (BBKNN) and ‘Mutual nearest neighbors correction’ (MNN) [31,32] and visualise individual cells on t-distributed stochastic neighbor embedding (tSNE). The BBKNN and MNN-correct approaches apply correction to neighbourhood graph and expression space respectively. The batch correction had minimal impact on resolving and overlapping similar cell types between the two atlases, compared to uncorrected data (Fig. S12A). Notably, the corrected batch effects were unique to each method on tSNE space. Performing SCENIC on uncorrected and two batch-corrected datasets, we find that individual regulon activities (RAS similarity) and regulon compositions (Jaccard coefficient) are highly correlated, indicating that batch effects have little effect on regulon activity (Fig. S12C-D). In summary, the pseudobulk approach accounts for technical and batch effects, robustly reports on regulon activities and leads to better classification of cell groups across individual and integrated atlas.

For an unbiased identification of concerted regulon activity across integrated atlas, we perform cell-to-cell correlation on RAS (Fig. 2A). We observe 3 major clusters with the largest cluster 1 composed of Immune and Stem cell groups from all atlases (Fig 1C-D). The cluster 2 is composed of Epithelial and Stem cell group, exclusively from MCA dataset, with several sub clusters within. The distinct MCA sub-clustering is expected, owing to increased sampling of tissues and single-cells (150,889 MCA vs 93,753 TM; S6A, C). The third cluster is composed of Stem and Specialised cell groups from all atlases. We observe several smaller clusters composed of individual cell-types highlighting their distinct classification based on specific regulon activity (Fig 2A). Next, we performed cell-to-cell correlation within individual atlases to identify clusters composed of shared and individual cell groups, highlighting the diversity of cell-types captured within each atlas. Consistent with integrated atlas, the shared clusters include ‘Stem and Specialised’, ‘Immune and Stem’, ‘Basal and Endothelial’, and are quite distinct from individual cell group clusters (Immune, Stem, Epithelial, Basal etc.,) in each atlas (Fig. S13A). To further investigate shared regulatory activity across individual atlases, we performed pairwise comparison and observed strong correlation between both shared and individual cell groups (Fig. S13B). In summary, the shared and individual cell group clusters validate that regulon activities correspond to true regulation in matched cell-types.

To highlight the regulon crosstalk and regulation across integrated cell atlas, we performed regulon-to-regulon correlation using Connection Specificity Index (CSI; Supplementary methods) [21,33]. The CSI is a context dependent graph metric that ranks the regulon significance based on similarity and specificity of interaction partners, thereby mitigating the effects of non-specific interactions. Correlating across integrated atlas using CSI, we identify 174 regulons across 5 distinct modules and sub-modules within (Fig 2B). For broad assessment of module features, we perform Gene Ontology (GO) using all genes within regulon modules and pathway analysis on regulons (Fig. S14A-B). The module 1 consists of 19 regulons (7165 genes) involved in various cellular processes including differentiation, metabolism and signal transduction predominantly in immune pathways (Fig. S14A-B). The module 2 consists of broad acting critical transcription factors Gata3 and Klf16 (605 genes) that regulate multitude of celltypes. Module 3 is composed of 66 regulons (8655 genes) involved in cellular differentiation, organogenesis (including Hox and AP1 family TFs) and with significant enrichment for signal transduction pathways (Fig. S14B). Module 4 consisted of 74 regulons (9286 genes) composed of core transcriptional activators with cell cycle and messenger RNA roles (E2F, SP, IRF family TFs), across both GO and pathway analysis. Lastly, cluster 5 is composed of 13 regulons (3344 genes) involved in broad roles in development, tissue and cellular organisation. Next, we compared whether regulon modules are distinguished based on CSI scores within individual atlases. The larger regulon modules (Module 3 and 4) are clearly separated within individual atlases, highlighting their roles in global regulation across multiple cell groups (Fig. S15A-C). The smaller modules (Module 1, 2 and 5) highlight tissue-specific regulation of different cell groups in both integrated and individual atlases. For example, the module 1 regulon Mafb regulates a subset of myeloid immune cells from microglia (Fig. 2B, Brain microglia in Fig S8A) [34], while the module 5 regulon Sox2 regulates Stem and Immune group (Fig. 2B, S9A).

To investigate regulon crosstalk within and between modules across the integrated atlas, we devised an undirected regulon network considering the most interacting regulons with stringent CSI association (CSI>0.7; Fig 2C). As expected from CSI correlation matrix (Fig. 2B), the regulons within modules have higher connections than across modules implying concerted regulation in cell-types across integrated atlas. We also assess several network features to determine regulon importances for individual modules as well as regulon network. Notably, the smaller modules (1, 2 and 5) bridge the nodes between larger modules (3 and 4) within the network. Within individual atlases, we find that the global regulon network is largely retained irrespective of regulon composition differences within atlas (Fig. S15D-F). We highlight regulons with important regulatory roles in reference cell types within individual atlases (Fig. S16). Assessing the several network features across the integrated network, we find Cebpd (module 1), Gata3 (module 2) and Hdac2 (module 5) are the key bridge nodes (betweenness centrality) for shortest path through the network. The top intra- and inter-module regulons have highly correlated network features (Degree, closeness and Eigencentrality with regulon composition; Fig 2C bottom). The network features across integrated atlas are detailed in Table S4.

To further validate the modules across integrated regulon network, we perform several in-silico comparisons. Firstly, our framework includes RCisTarget (as a part of SCENIC) for defining regulons i.e. TFs and direct target genes. RCisTarget crossmatches identified regulons with known and annotated TF-target databases, prunes indirect co-expressed targets and enables scoring of TF-TF and TF-target relationships. Consequently, all direct targets of a given regulon harbour the regulon motif at respective promoters. Additionally, we expect and observe many regulons within individual modules to share overlapping motifs (Motif correlation in Fig. S17A). We also report a few representative examples of regulons and their motifs within individual modules (Fig. S17A). We next assessed whether regulons crosstalk across the integrated network are mediated through protein-protein interactions (PPi). Comparing and overlaying the annotated PPi from STRING [35], we validate 57% of regulon network connections (Fig. S17B). Since each STRING annotated PPi is assigned a combined score (a measure of confidence), we compared our regulon network with the STRING combined score (in 20% bins; Fig. S17C). We find that the regulon network connections have the highest STRING combined score. Additionally, we also observe a strong positive relationship between regulon CSI and STRING combined score, validating the regulon network interactions from experimental evidence (Fig. S17D, Table S5). Lastly, we compared our regulon network for essential genes in the Online Gene Essentiality database (OGEE) [36]. We observe 109 essential genes (70%) in our regulon network with strong representation across all modules (Fig. S17D, Table S5), further highlighting the regulon importance across integrated network.

Next, we focussed on regulons with differential composition that drive individual cell-types (Fig. 3A, S16). The regulon Cebpe consists of 1342 unique genes (TM-10x: 332, TM-SS2: 531 and MCA: 479 genes) with 189 common and direct targets. The Cebpe activity is highly specified in granulocyte and monocytes, consistent with its known role in lineage determination (Fig. 3A) [37]. The Irf8 is a master regulator of monocytes, dendritic cells and is important for both adaptive and innate immunity [38]. We observed 641 shared targets and specific activity in monocytes and macrophages (Fig 3B). We find that regulons with few shared direct targets across cell atlases have specific and consistent activity. The Lef1 and Hoxb7 regulons have fewer overall targets genes, only 5 shared targets between cell atlases, but with specific activity in T-cells [39] and kidney epithelial cells [40] respectively. Several global and cell-type specific regulons with differential compositions are presented in Fig. S18-20.

To further validate our regulatory framework for atlas-scale analysis, we performed GRN inference using an alternative method ‘bigSCale2’ considering TM-10x atlas [41]. The ‘bigSCale2’ approach uses expression correlation to calculate regulatory network and does not distinguish between direct and indirect TF-targets. Comparing the two methods, we find 117 regulons (67%) co-identified by both methods, while 57 regulons (33% and direct targets within) exclusively captured in our SCENIC framework (Fig. S21A). Computing the Jaccard index, we find only 95 regulons with composition similarity between both GRN inference methods (Fig. S21B). In summary, the SCENIC framework robustly identifies regulons and their direct targets for atlas-scale analysis.

Lastly, we assess the functional importance of regulon activity by investigating mixed-lineage transitions during myeloid cell-fate determination using scRNA-seq [42]. The Irf8 regulon and its regulatory interactions are critical for monopoiesis and have a reciprocal dynamics with Gfi1 driven granulocyte specification [42]. We analyse granulocytic and monocytic specification in wildtype and Irf8-/- progenitors using scRNA-seq data (Fig. S22A), infer regulons and score regulon activity in single-cells (Fig. S22B). Both scRNA-seq expression counts and regulon activities separate different cell-types and capture the shift in Irf8-/- cells towards granulocyte lineage (Fig. S22A-B). Comparing the Irf8 regulon across monocytes, granulocytes and Irf8-/- cells, we observe preferentially high composition of direct targets in Monocytes (542 genes) over granulocytes (148 genes), consistent with cell-fate roles (Fig. S22E) [42]. Notably, the Irf8 regulon is significantly perturbed both in composition (direct target genes) and activity (target regulation) across Irf8-/- cells (Fig. S22D-E), highlighting the functional importance of regulon activity in mediating cell states and cell-types.

## Discussion

As major tissue, organ and organism expression atlases are generated [43,44], it is critical to also decipher mechanistic gene regulatory programs for refining functional cell-state and cell-type definitions. Here, we highlight a computational approach to infer regulons and link their specific cell-type activity with functional roles, from integrated scRNA-seq cell atlases. To resolve the author assigned cell-type labels across cell atlases, we standardise and categorise single-cells into broadly defined 7 cell groups and reference cell types. In our study, we project and map cell groups across different cell atlases, which indeed diminishes the resolution of individual celltypes. However, it allows us to converge and group similar cell-types together irrespective of differences in cell atlases including tissue sampling, scRNA-seq platform and sequencing depth. For regulon identification, inference and scoring, we used GRNBoost, RcisTarget and AUCell (as in [10], however, alternative inference methods have been proposed with improvements for both directed and undirected networks [9,12,45,46]. A fundamental caveat of recent GRN methods is the requirement of *a priori* pseudotime or cellular trajectory, which makes them incompatible for atlas-scale analysis. Our integrative analysis on three mouse cell atlases uncovers global regulon modules operating on multiple cells types, as well as specialised regulons critical for cell-type definition and identity. Through a variety of in-silico comparisons, we highlight the robustness of pseudobulk cells in effectively classifying cell groups and highlight regulatory crosstalk. The global regulon network is recapitulated in individual cell atlases using both single- and pseudobulk cells, validating the regulatory crosstalk in individual cell groups. The functional consequence of regulon composition and activity is highlighted during the lineage transition from monocytes to granulocytes in Irf8-/- cells. Our integrated computational atlas with standardised classification of cell groups, global and cell-type specific regulons across three mouse cell atlases presents a valuable resource for single-cell community.

## Supplementary Material

1. **Supplementary methods and text** - Supplementary methods and text (SupplementaryText.docx; 27kb)

2. **Supplementary Table 1** - The contingency table with pairwise cell-type mapping between atlases (SuppTable1.xlsx; 171kb)

3. **Supplementary Table 2** - The regulon composition and modules in the integrated cell atlas and across individual cell atlases (SuppTable2.xlsx; 1809kb)

4. **Supplementary Table 3** - The mean RAS score for individual pseudobulk cell types across integrated and individual atlases (SuppTable3.xlsx; 575kb)

5. **Supplementary Table 4** - The summary of different network features for regulons across modules in the integrated atlas (SuppTable4.xlsx; 829kb)

6. **Supplementary Table 5** – Comparison of integrated regulon network with STRING and OGEE databases (SuppTable5.xlsx; 1016kb)

## Declarations

### Data Availability

The supplementary document contains the full data sources analysed in the current study. The Jupyter notebooks detailing all the analysis steps can be found here: https://github.com/Nataraianlab/Single-cell-regulatory-network.

## Acknowledgements

The authors thank Lars Grontved and other Functional Genomics Unit members for helpful discussions, and comments on the manuscript. The authors also thank DeIC National HPC Centre (ABACUS 2.0) at SDU, Denmark for computational resources.

## Author contributions

KNN designed and supervised the project. AFM performed the analysis with help from KNN.

KNN wrote the manuscript. Both authors reviewed and approved the manuscript.

