## Supplementary Figures for "Predicting gene regulatory networks from cell atlases"

**Supplementary Figure 1: Detailed workflow of atlas-scale GRN analysis.**
To effectively integrate single-cell annotations across three atlases, we first manually devised 7 reference cell groups, chose TM-10x as a common reference and assigned the 55 cell types to 7 cell groups respectively. Next, we used scMAP to link each atlas (TM-SS2 and MCA) to the reference and built an integrated mouse atlas with common vocabulary for all single-cells. Using a stringent feature selection cutoff, we performed gene regulatory network inference using SCENIC. This briefly includes TF and TF-target identification from single-cell expression matrices (GRNBoost), cross-validation of TF and its direct targets (i.e. Regulons) using annotated motif databases and pruning away indirect, co-expressed genes (RCisTargets) and lastly scoring the regulon activity (RAS: regulon activity score) within each single-cell. We applied the framework to integrated and individual atlases to *(i)* classify individual and pseudobulk cells based global regulon activity (UMAP), *(ii)* classify cells based on shared and distinct regulon activity (cell-to-cell correlation), *(iii)* identify consensus and cell group specific regulons (regulon-to-regulon correlation) and *(iv)* build an atlas-scale regulon activity network.

**Supplementary Figure 2: Cell type and cell group relationships**
**A.** Simplified mapping and projection of 55 reference cell types to 7 *cell groups.*
**B.** scMAP visualisation of TM-10x cell atlas (reference onto itself) with 55 reference cell-types from 12 tissues.

**Supplementary Figure 3: Mapping cell type annotations across individual cell atlases**
**A.** scMAP projection of TM-SS2 with 81 unique author assigned cell-type labels from 18 tissues to 55 reference cell types.
**B.** scMAP projection of MCA with 732 unique author assigned cell-type labels from 38 tissues to 55 reference cell types.

**Supplementary Figure 4: Mapping cell type annotations across integrated cell atlas**
**A.** scMAP projection of integrated mouse atlas with 831 unique author assigned cell-type labels from 50 tissues to 55 reference cell types.

**Supplementary Figure 5: Examples of cell type annotations within individual tissues across integrated cell atlas.**
**A.** Projection of Liver cell types to reference cell groups across integrated cell atlases
**B.** Projection of Spleen cell types to reference cell groups across integrated cell atlases

**Supplementary Figure 6: Technical differences and biases in mouse cell atlases**
**A.** Number of cells (first row), library size (i.e. sequencing depth; second row) and number of genes detected (third row) for individual and integrated atlas. Each point represents a single-cell and the metrics are stratified and coloured based on 7 cell groups and 55 cell types with the median numbers reported next to box plots.

**B.** Relationship between mean expression and dropout rate (percentage) for each atlas. TM-SS2 was more deeply sequenced and utilises full length Smart-seq2 protocol, while TM-10x and MCA apply 3' end scRNA-seq protocols.
**C.** Simplified table highlighting number of single- and pseudobulk cells, stratified by 7 cell groups across integrated cell atlas.
**D.** Pie charts showing proportion of single- (top) and pseudobulk (bottom) cells across the 7 cell groups in individual and integrated cell atlas. Note: proportions are conserved between single- and pseudobulk cells.

**Supplementary Figure 7: Technical differences and biases in cell atlases stratified by cell groups and reference cell types**
**A.** Number of cells (first row), library size (i.e. sequencing depth; second row) and number of genes detected (third row) for individual and integrated atlas stratified for each cell group and reference cell types. Although the 7 cell groups (and 55 reference cell groups within) are well represented from individual atlases; the Basal and Endothelial cells are under-represented in MCA.

**Supplementary Figure 8: Tissue profiled across integrated mouse atlas and examples of regulon in single- and pseudobulk cells**
**A.** UMAP embedding of single-cells across integrated atlas based on RAS (same as Fig 1C), coloured by unique tissues across all cell atlases.
**B.** UMAP embedding of pseudobulk across integrated atlas, based on RAS, coloured by unique tissues across all cell atlases.
**C.** Examples of tissue- and reference cell-type specific regulons (Pou2af1, Cebpa, Gli1, E2f7 and E2f4), coloured by RAS.
**D.** Examples of general and cell-type specific regulons (Atf3, Elk3, Foxc1, Trp53), coloured by RAS.

**Supplementary Figure 9: Cell type and tissue specific regulon across individual atlases.**
**A.** UMAP embedding of single cells in individual atlases coloured by reference cell types (top), RAS for immune Irf8 (middle) and cell type specific regulons (bottom: Sox17, Sox2 and Pparg)
**B.** UMAP embeddings of pseudobulk cells in individual atlases coloured by reference cell types (top), RAS of tissue-specific regulons (middle: Snai3 and bottom: Tcf7).

**Supplementary Figure 10: Individual atlases stratified by tissues.**
UMAP embedding of (**A**) single-cells and (**B**) pseudobulk cells across individual cell atlas, coloured by unique tissue types.

**Supplementary Figure 11: RAS comparison between single- and pseudobulk cells across integrated and individual atlases**
**A.** Distribution of Gini coefficients per regulon in pseudobulk and single-cells across integrated atlas, stratified by individual cell groups.
**B.** Silhouette score comparing clustering between single- and pseudobulk cells across individual and integrated atlas.

**C.** RAS correlation between single and pseudobulk cells in all cells and stratified by individual cell groups. Error bars represent the standard deviation across single- and pseudobulk cells. The individual cell group correlation is significantly improved compared to global, which further validates our classification of 7 cell groups.

**Supplementary Figure 12: Impact of batch effect correction on regulon inference**
**A.** UMAP embedding of pseudobulk cells from Spleen in both TM-10x and TM-SS2 atlases, considering either uncorrected or two batch corrected expression space (BBKNN and MNN-correct). The pseudobulk cells are coloured by cell atlas (top) and cell groups (bottom).
Note: Both batch correction methods slightly improve the overlap of reference cell types compared to uncorrected UMAP. However, the clusters from BBKNN and MNN-correct don't overlap with each other and introduce additional discrepancies
**B.** Pairwise correlation of individual regulons (based on RAS) from both batch correction methods compared to uncorrected data. Each dot represents a regulon identified in all 3 SCENIC runs (uncorrected, BBKNN and MNN-correct). The shaded area represents the 95% confidence interval from the linear regression line.
**C.** Regulon composition similarity computed from pairwise Jaccard index between batch corrected (BBKNN and MNN-correct) to uncorrected data. The shaded area represents the 95% confidence interval from the linear regression line.

**Supplementary Figure 13: Cell-type correlation across individual cell atlases**
**A.** Spearman correlation map of reference cell types within each cell atlas using pseudobulk cells. The first column highlights reference cell types. The cluster comparison between individual atlases and integrated atlases is presented in Fig. 1E.
**B.** Pairwise correlation between individual cell atlases ('TM-10x vs TM-SS2', 'TM-10x vs MCA' and 'TM-SS2 vs MCA'). The same colour scheme is used in **A** and **B** to indicate 55 reference cell types

**Supplementary Figure 14: Regulon module features**
**A.** Significant Gene Ontology (GO; biological processes) terms for each module. All regulons and direct targets within modules are used for GO analysis and are listed under the module (X-axis). The gene ratio highlights percentage of total GO term genes identified as enriched within the module. The gene list considering regulons alone (without direct targets) was too sparse to produce significant GO terms per module.
**B.** Reactome pathway analysis using regulons within modules. Only regulons within modules are used for Pathway analysis and are listed under the module (X-axis). Only significant (qval<0.05) and enriched edges were considered for quantification and highlighted as tick sizes.

**Supplementary Figure 15: Regulon correlation across individual cell atlases**
**A-C.** Connection specific index (CSI) matrix for each cell atlas. The identified regulons are coloured as in Fig 2B (5 modules), indicating co-regulatory and distinct roles across different cell types and tissues. Regulons identified in individual atlases but not in integrated atlas are marked in grey.

**D-F.** Regulon network for each cell atlas. Each regulon is represented as a node, nodes are connected if CSI scores > 0.7 and coloured as in Fig 2C (5 modules). Both shared and unique regulon architecture are conserved in each cell atlas. Similar to integrated network (Fig. 2C), the larger modules (3 and 4) are bridged by smaller regulons (1, 2 and 5) within each cell atlas. The CSI for two nodes A and B is calculated by:

$$CSI_{AB} = 1 - \frac{\#nodes\ connected\ to\ A\ or\ B\ with\ PCC \geq PCC_{AB} - 0.05}{n_y}$$

The Pearson correlation coefficient (PCC) is the interactional correlation between A and B.

**Supplementary Figure 16: Regulon activity in reference cell types across individual cell atlases.**
**A-C.** Individual regulon activities highlighted in regulon-by-cell type matrix for (**A**) TM-10x, (**B**) TM-SS2, and (**C**) MCA.

**Supplementary Figure 17: Validation of regulon network**
**A.** Top: Motif correlation between individual regulons within each module. The rows and columns indicate individual motif sequences of different lengths. Bottom: representative examples of TFs and their enriched motifs for each regulon module.
**B.** Annotated protein-protein interactions (PPi) from STRING overlaid on integrated regulon network. STRING contains all regulons (nodes), and only STRING validated interactions (black edges) are highlighted in the regulon network. 57% of regulon network edges are validated by STRING.
**C.** Distribution of the STRING validated interactions captured in regulon network, plotted across 20 percentile combined score bins (x-axis). The number of regulon network links are listed above individual. The combined score is measure of confidence of STRING PPi.
**D.** Correlation between regulon connection specificity index (CSI) and STRING confidence score. The error bars represent the 95% confidence interval. Red line indicates the CSI threshold used to construct regulon network.
**E.** Regulon network overlaid with experimentally validated and essential genes (OGEE essentiality status). The enlarged nodes represent essential genes, while diminished nodes are annotated as non-essential. The regulons absent in OGEE are shown in 'grey'.

**Supplementary Figure 18, 19 and 20: Example for individual regulons composition and activity in reference cell types across cell atlases.**
Fig. S14. **(A)** Pou2af1, (**B**) Eomes and (**C**) Tcf7.
Fig. S15. **(A)** Hsf1, (**B**) Etv3 and (**C**) Mafb.
Fig. S16. **(A)** Irf5, (**B**) Irf9 and (**C**) Foxp1.

**Supplementary Figure 21: Comparing different GRN methods for atlas-scale analysis**
**A.** Overlap of regulons and target genes identified in this study (SCENIC: GRNBoost and RCisTarget) and repeating analysis with published GRN method (bigSCale2; Ioconno et al 2015) for TM-10x mouse atlas.
**B.** Jaccard index highlighting the overlap between regulon composition inferred on TM-10x atlas by our framework and repeating analysis with bigSCale2.

**Supplementary Figure 22: Functional importance of regulon activity during myeloid differentiation**

**A.** UMAP embedding based on scRNA-seq expression from wildtype (multiple cell types) and Irf8 KO (or Irf8-/-) single-cells.

**B.** UMAP embedding based on RAS from wildtype (multiple cell types) and Irf8 KO single-cells. The regulon space captures cell types differences as with expression space.

**C.** Single-cell Irf8 expression in myeloid progenitors from wildtype (incl. monocytes, granulocytes) and Irf8 knockout cells (Purple). The Irf8 KO cells have a gradient expression of high and low cells.

**D.** Irf8 regulon activity in myeloid progenitors from wildtype (incl. monocytes, granulocytes) and Irf8 KO cells (Purple). Note: the Irf8 KO cells have diminished RAS (compared to monocytes, consistent with change in their specification from monocyte to granulocytes.

For calculating Irf8 regulon activity in both wildtype and Irf8 KO cells, we repeated AUCell 50 times and use the averaged activity score.

**E.** UpSet plot of Irf8 regulon composition in wildtype monocytes, wildtype granulocytes and Irf8 KO cells. The Irf8 knockout cells alter and specify cell fate from monocytes to granulocytes, as highlighted by with drastically altered regulon composition in Irf8 KO cells.

Supplementary Figure 1

A. Detailed analyis workflow

Tabula Muris → 10x (~54,000 cells); Smart-seq2 (~38,000 cells)

Mouse cell Atlas → Microwell-seq (~150,000 cells)

**Map cell types to a common reference**
1. Manual annotation of atlas annotated cell types to 7 reference cell groups
2. TM-10x selected as reference. Assign 55 reference cell types to 7 reference cell groups
3. Use scMAP to map TM-SS2 and MCA author-annotated cell types to TM-10x (reference)
*(scMAP correlation threshold: 0.7)*

↓

**Feature selection for regulon inference** *(Cutoff : Genes expressed in at least 10% cells)*

↓

**Gene regulatory inference for integrated and each atlas (using SCENIC)**
1. GRNBoost/GENIE3: Finds co-expressed and correlated TF-gene pairs
2. RCisTargets: Prunes away co-expressed TF-gene pairs using annotated motif database
3. AUCell: Scores regulons (TF-gene pairs) within each single-cell

| **Integrated** | **TM-10x** | **TM-SS2** | **MCA** |
|---|---|---|---|
| Tissues: 50 | Tissues: 12 | Tissues: 18 | Tissues: 54 |
| Single-cells (SC): 244,642 | Single-cells (SC): 54,865 | Single-cells (SC): 38,888 | Single-cells (SC): 150,889 |
| Pseudobulk (PB): 4,550 | Pseudobulk (PB): 1,061 | Pseudobulk (PB): 703 | Pseudobulk (PB): 2,776 |
| Cell types: 831 | Cell types: 55 | Cell types: 81 | Cell types: 732 |
| Features: 17,138 | Features:19,369 | Features: 21,373 | Features: 34,967 |
| Regulons: 174 | Regulons: 222 | Regulons: 233 | Regulons: 233 |

**Regulon activities**
*(reference cell groups)*

**Cell-to-cell correlation**

**regulon-to-regulon correlation**

**Regulon activity network**

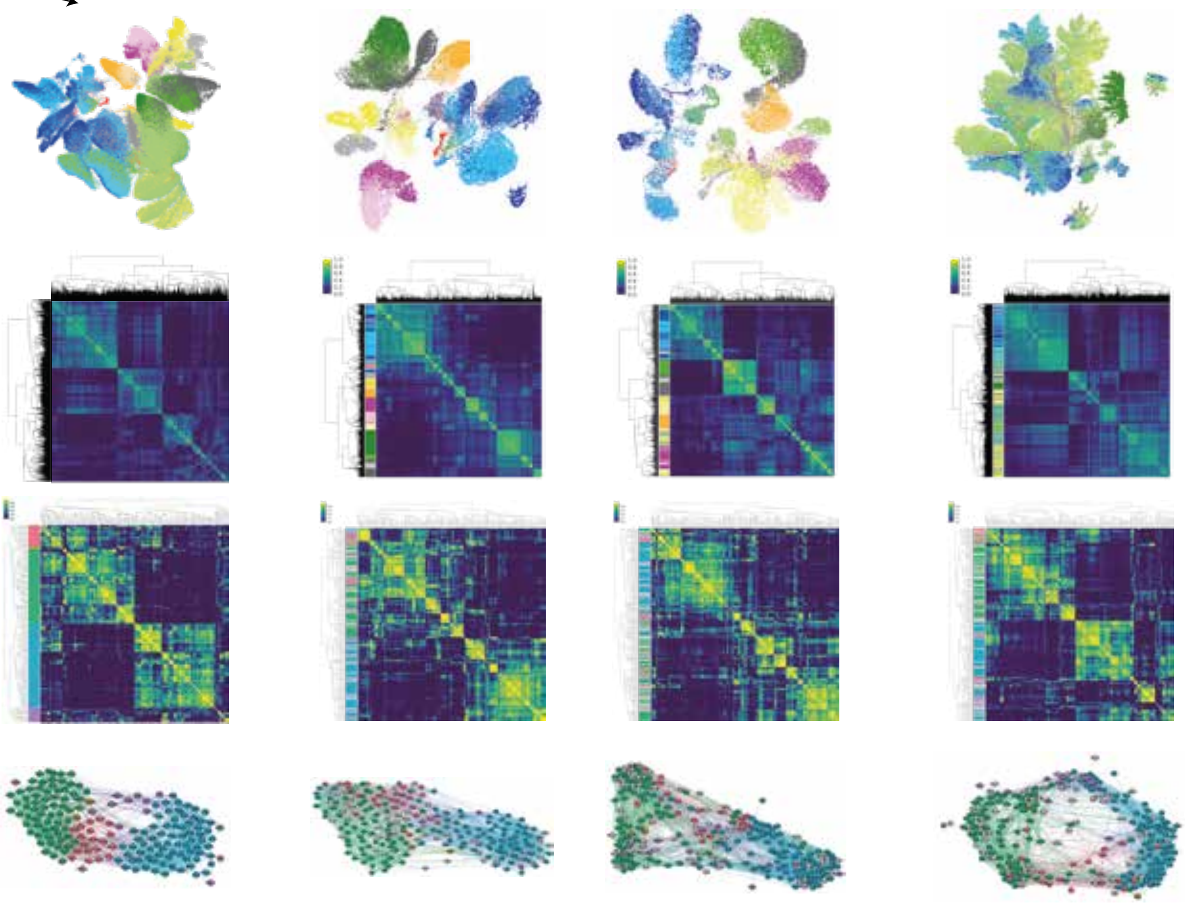

Supplementary Figure 2

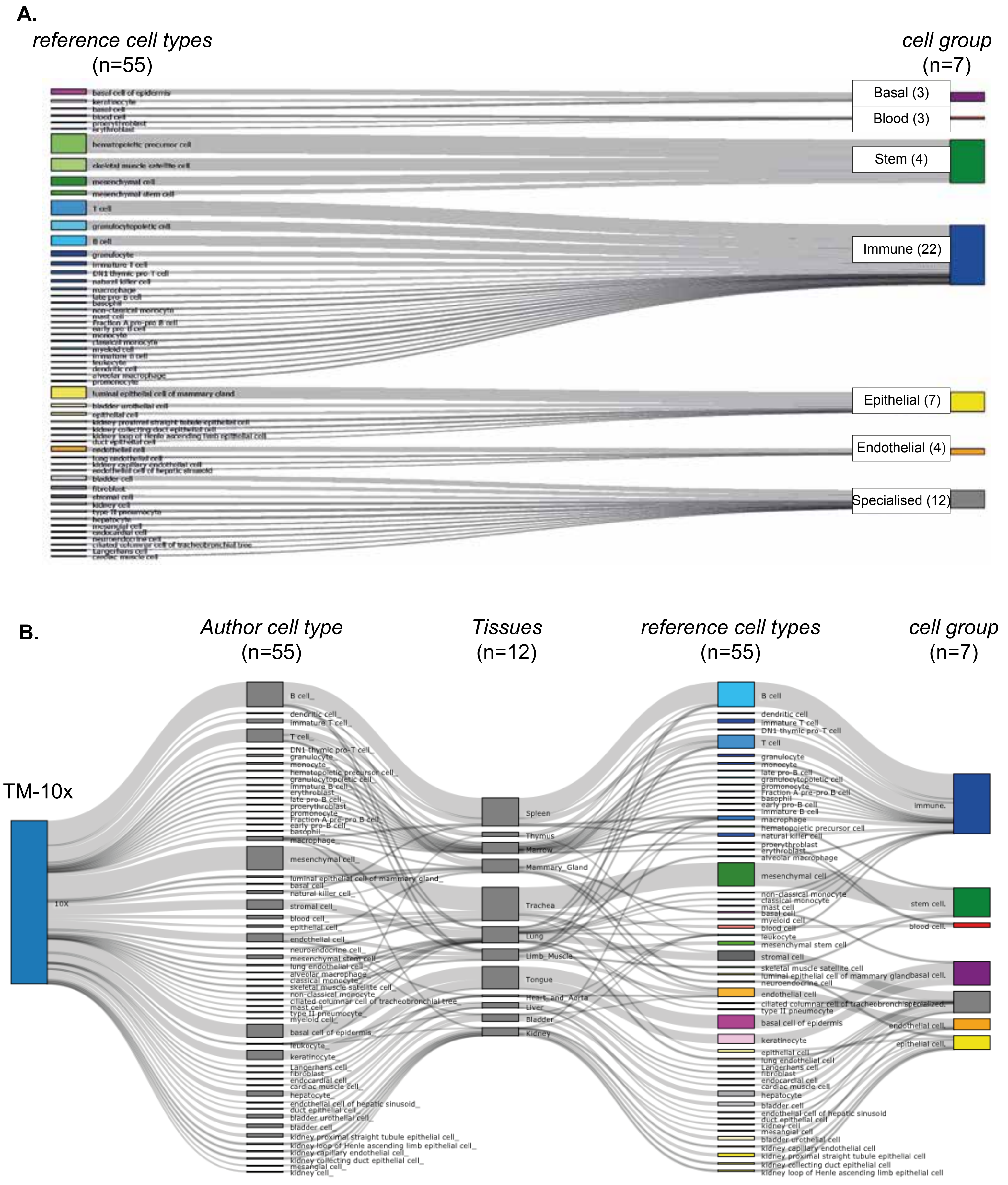

Supplementary Figure 3

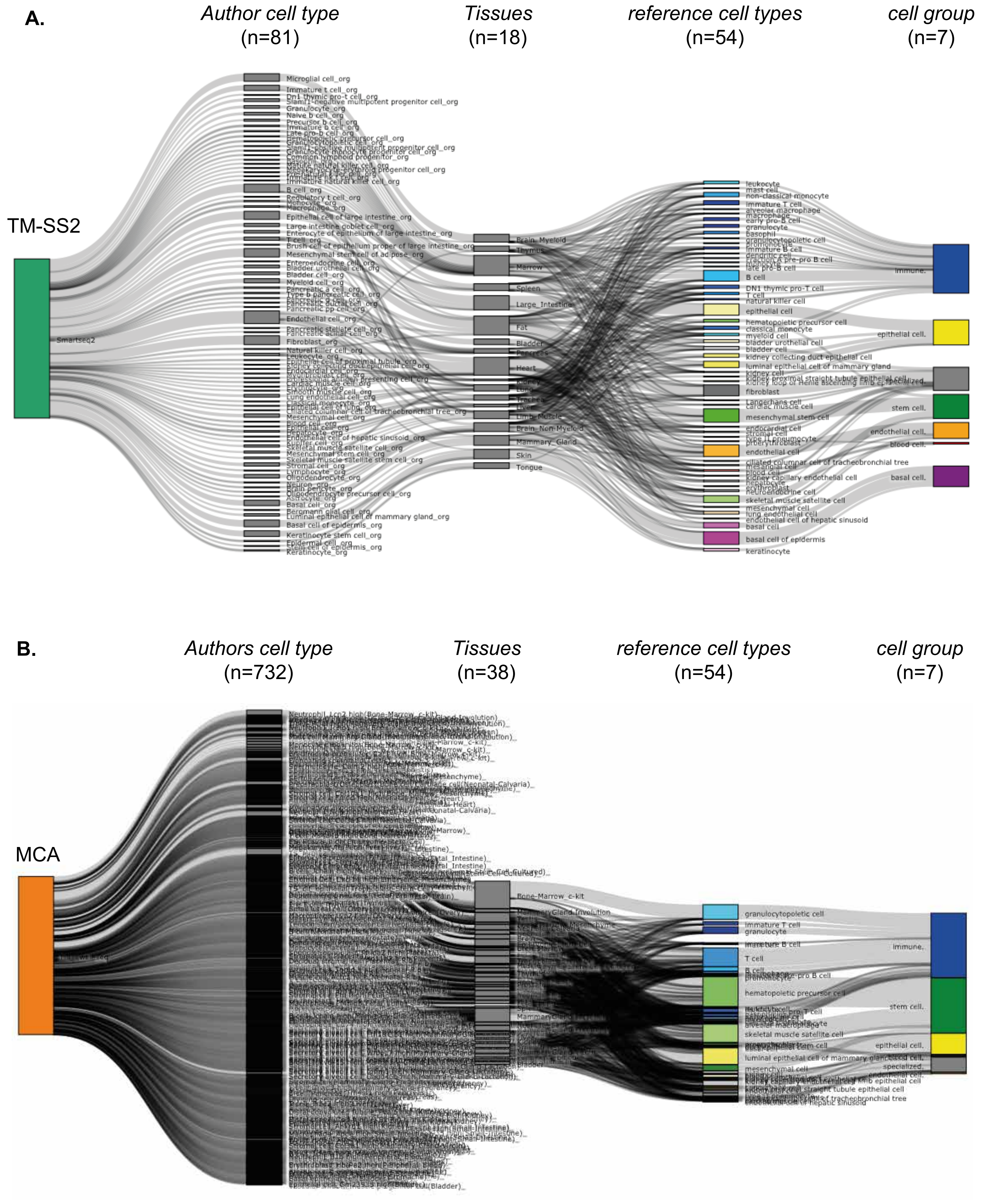

Supplementary Figure 4

**A.**

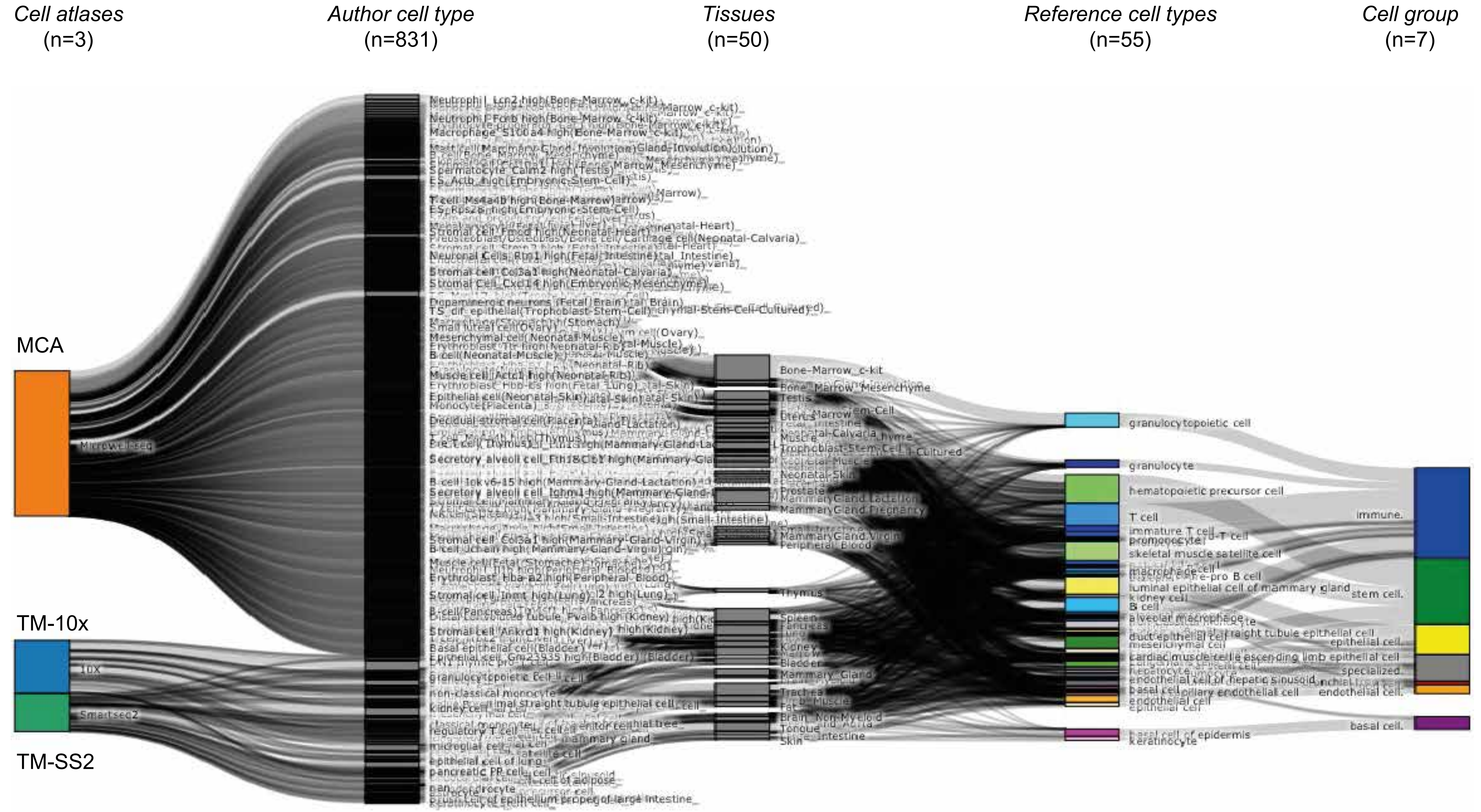

Supplementary Figure 5

A.

Cell atlases (n=3) | Author cell type (n=27) | reference cell types (n=36) | cell group (n=7)

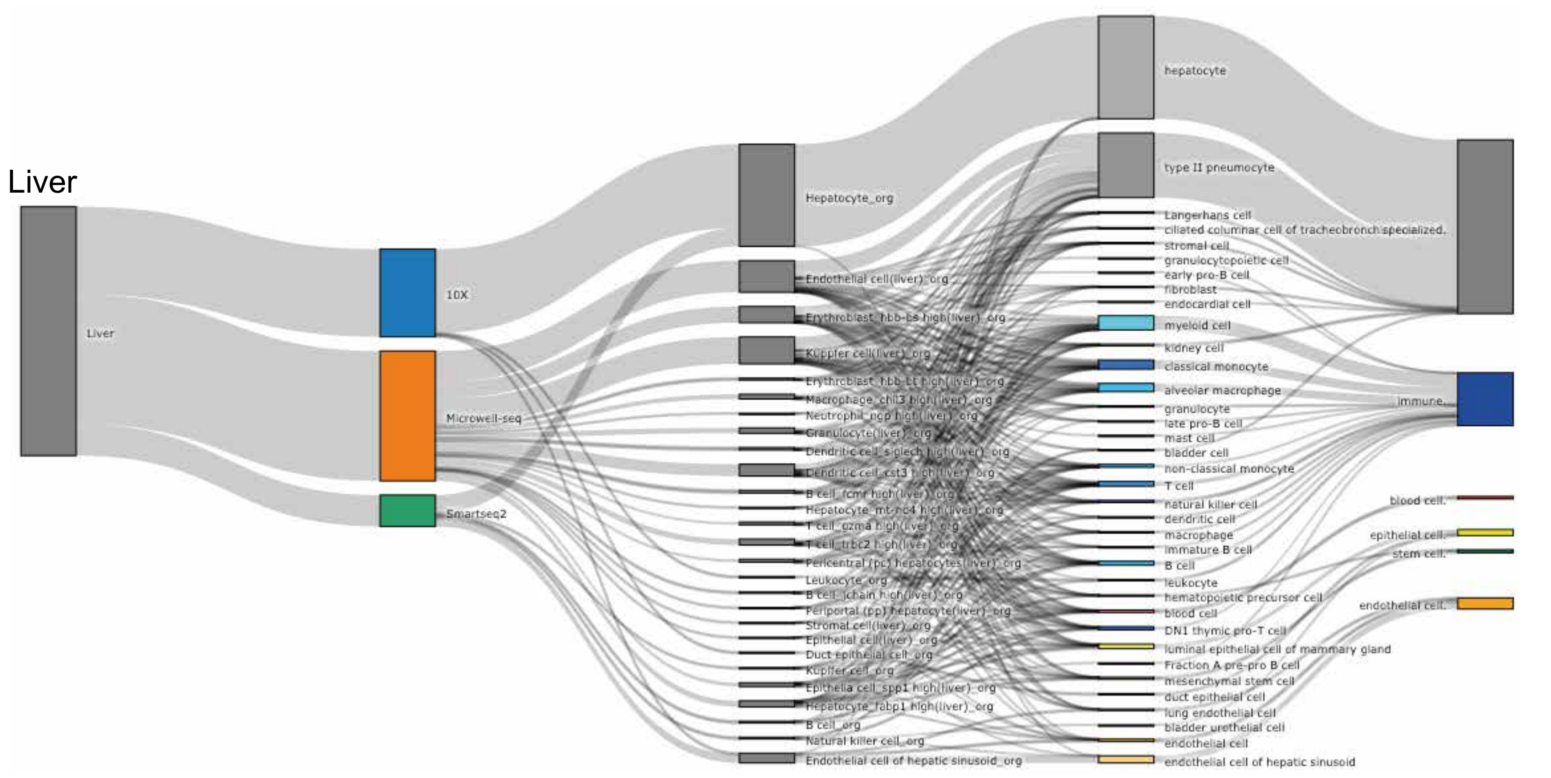


B.

Cell atlases (n=3) | Author cell type (n=16) | reference cell types (n=34) | cell group (n=7)

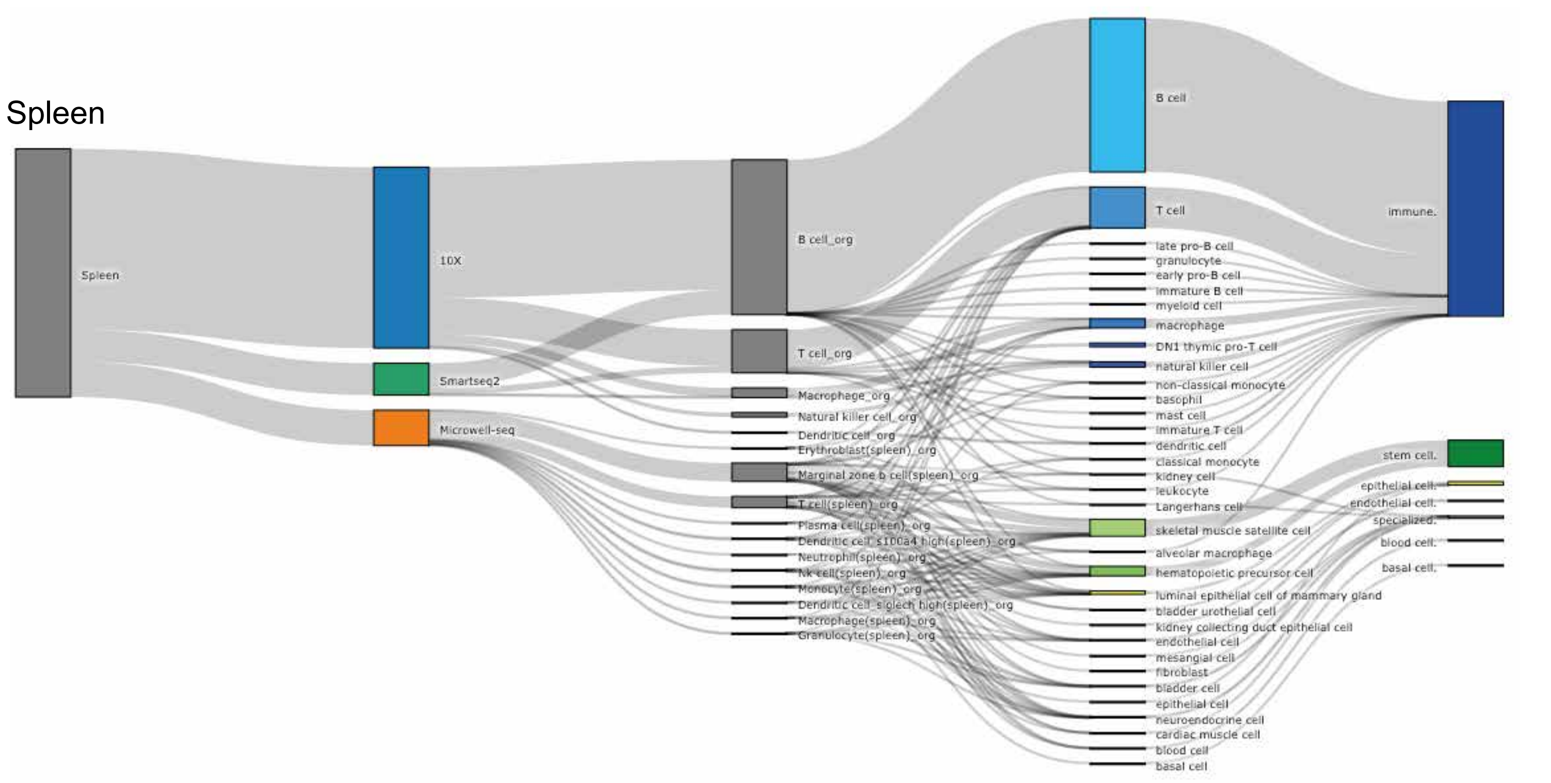

### Supplementary Figure 6

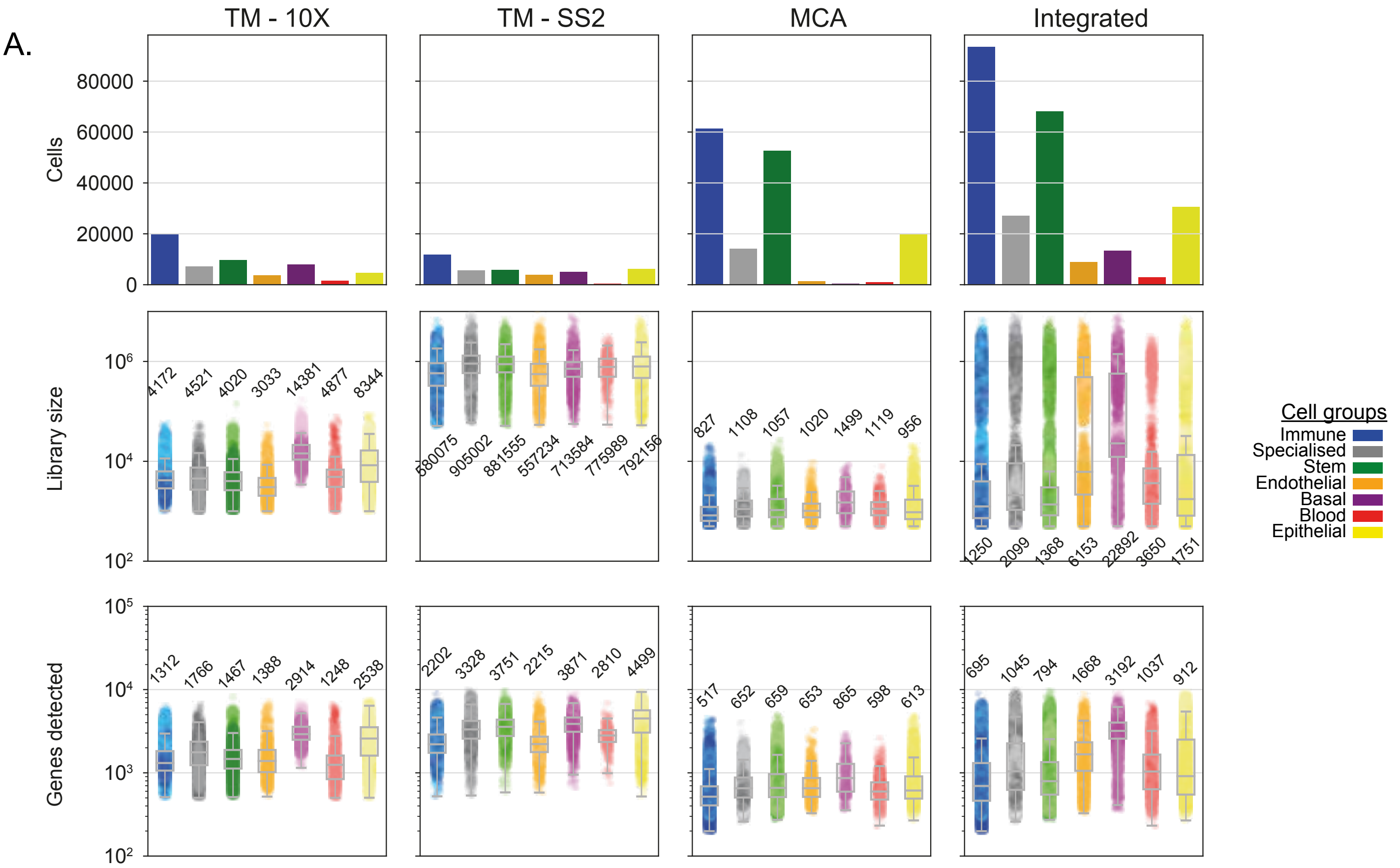


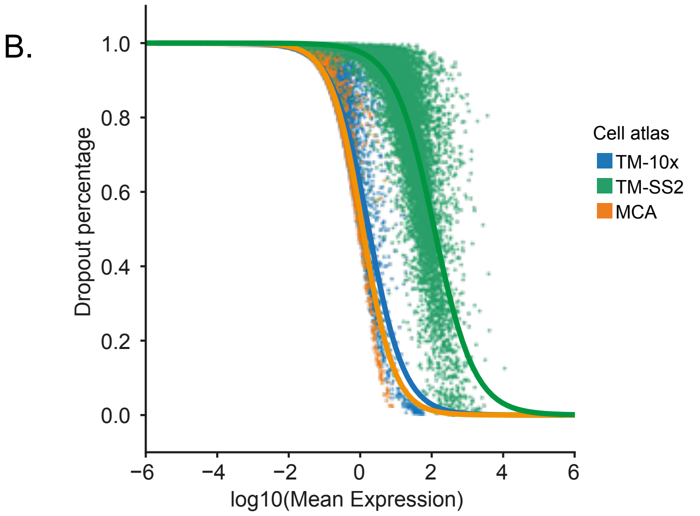


C.

#### Integrated mouse atlas cells

(by reference cell groups; also in Pie chat)

| | Single-cell | Pseudobulk |
|---|---|---|
| Basal | 13394 | 261 |
| Blood | 2925 | 46 |
| Endothelial | 8913 | 158 |
| Epithelial | 30660 | 584 |
| Immune | 93461 | 1702 |
| Specialised | 27078 | 479 |
| Stem | 68211 | 1310 |

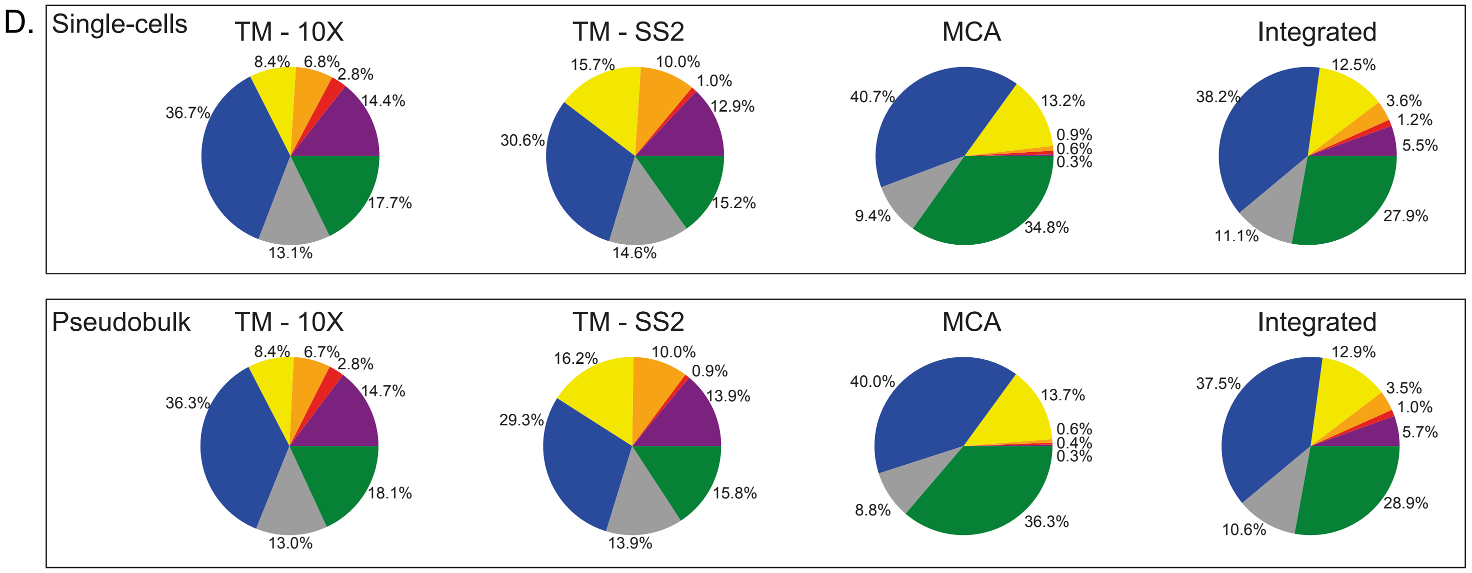

Supplementary Figure 7

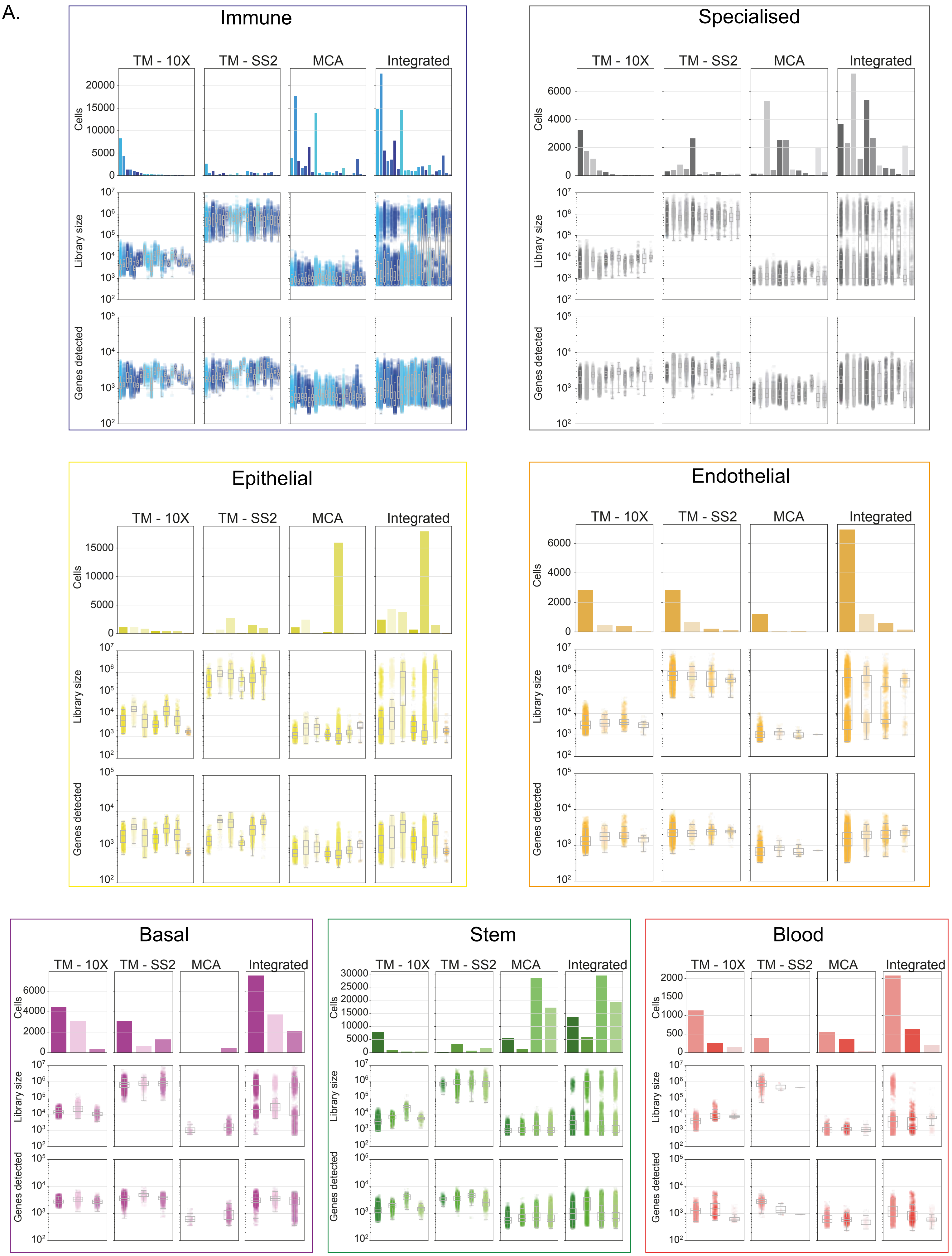

Supplementary Figure 8

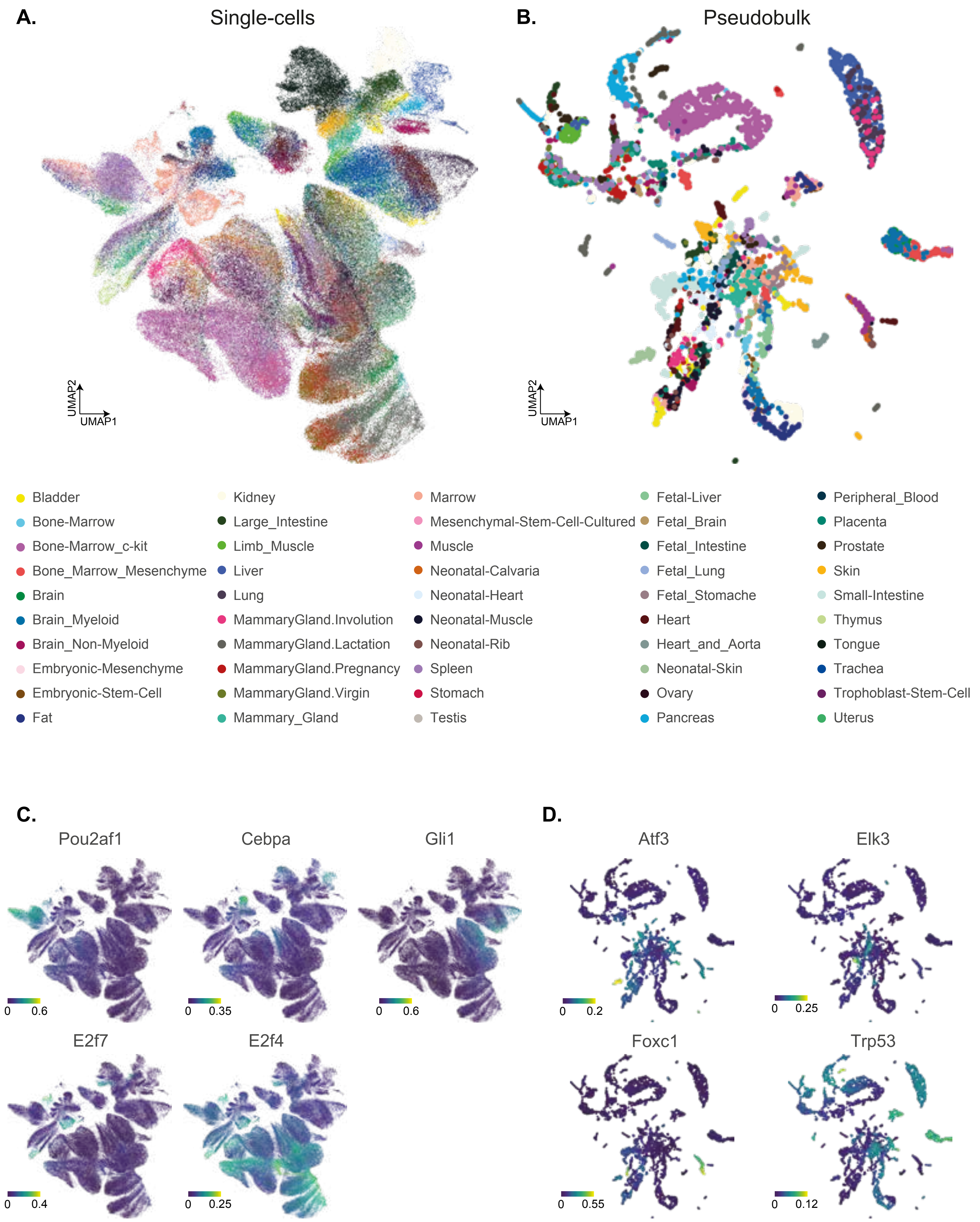

Supplementary Figure 9

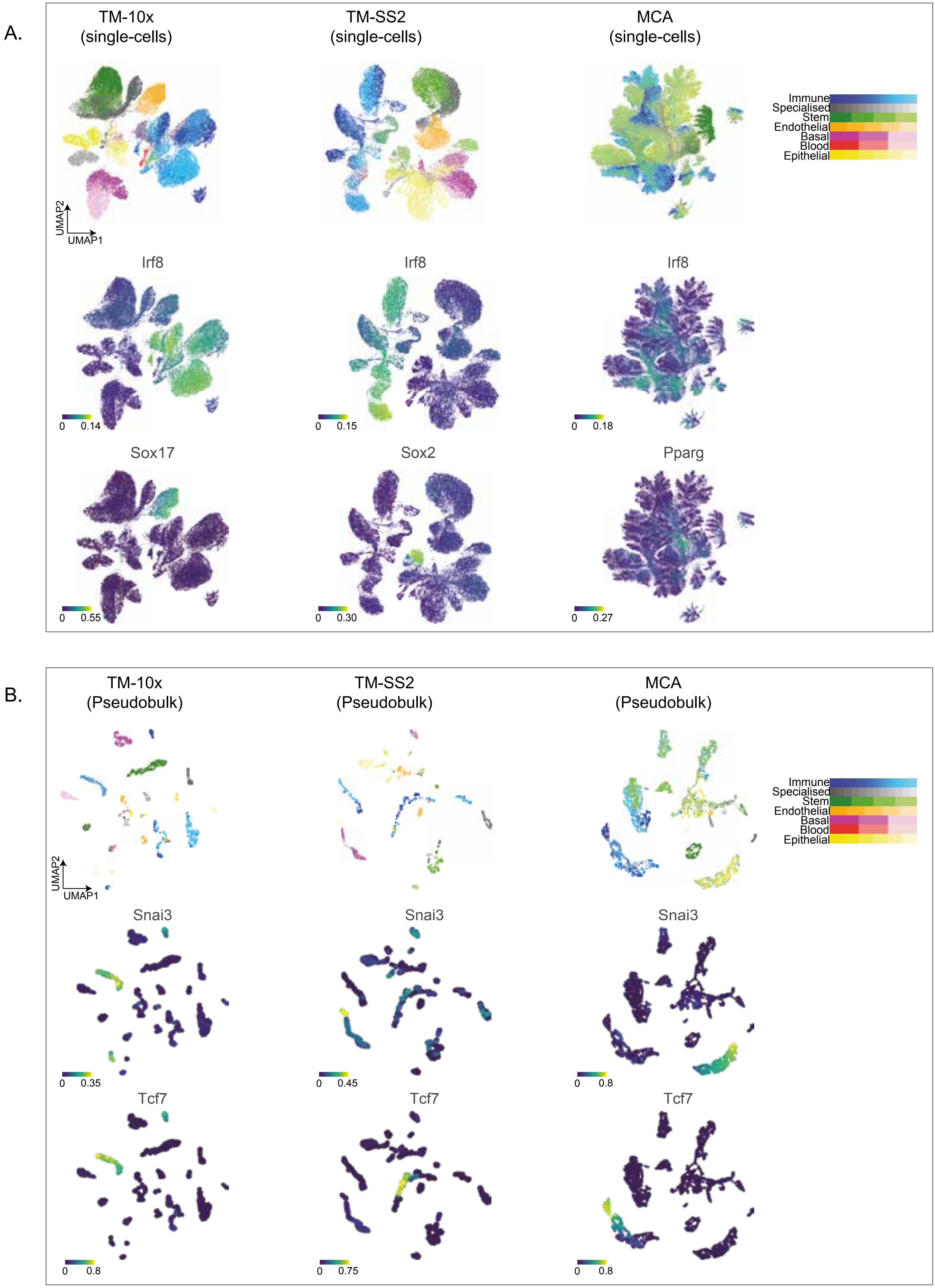

Supplementary Figure 10

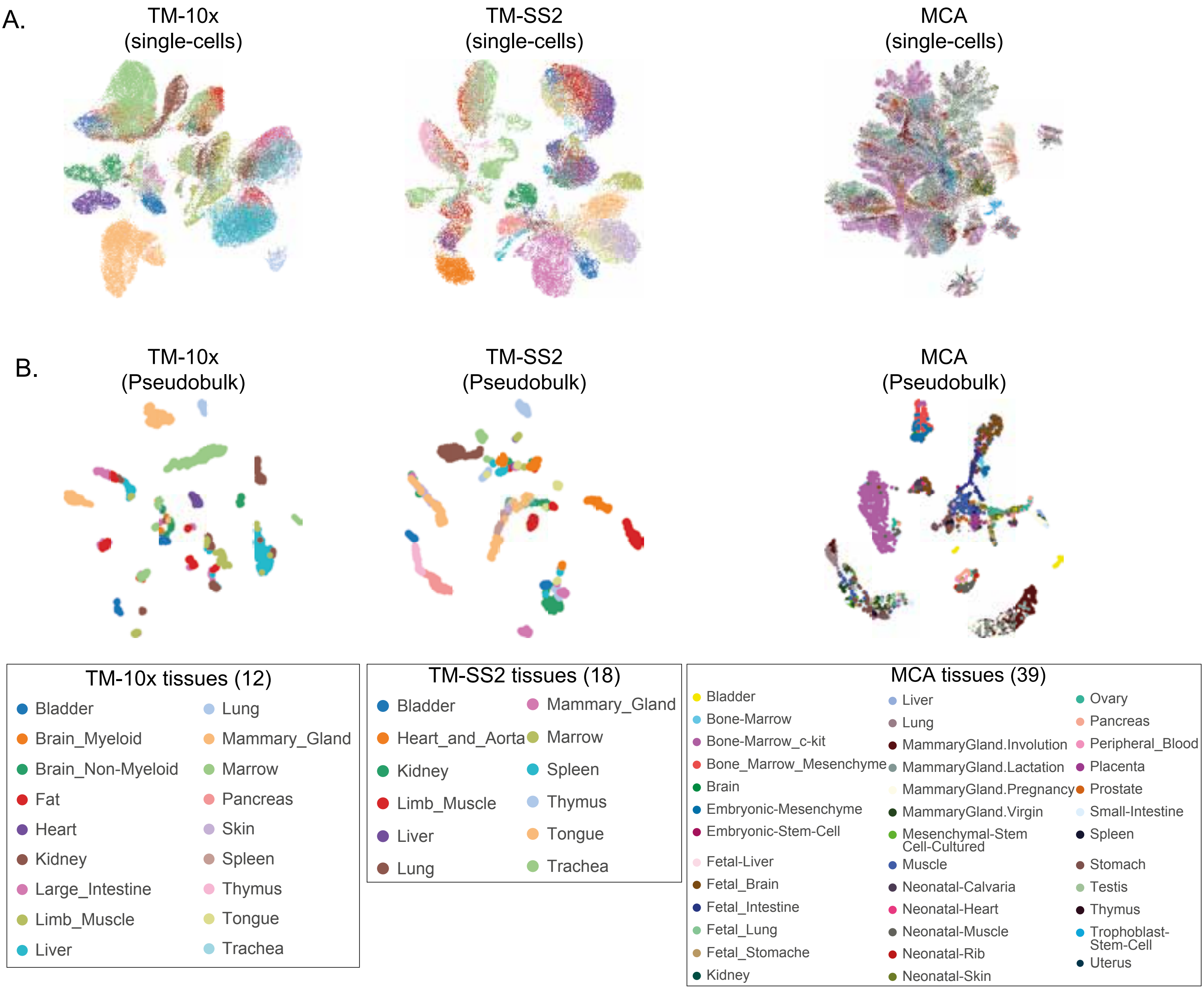

Supplementary Figure 11

**A.** PB vs SC comparisons

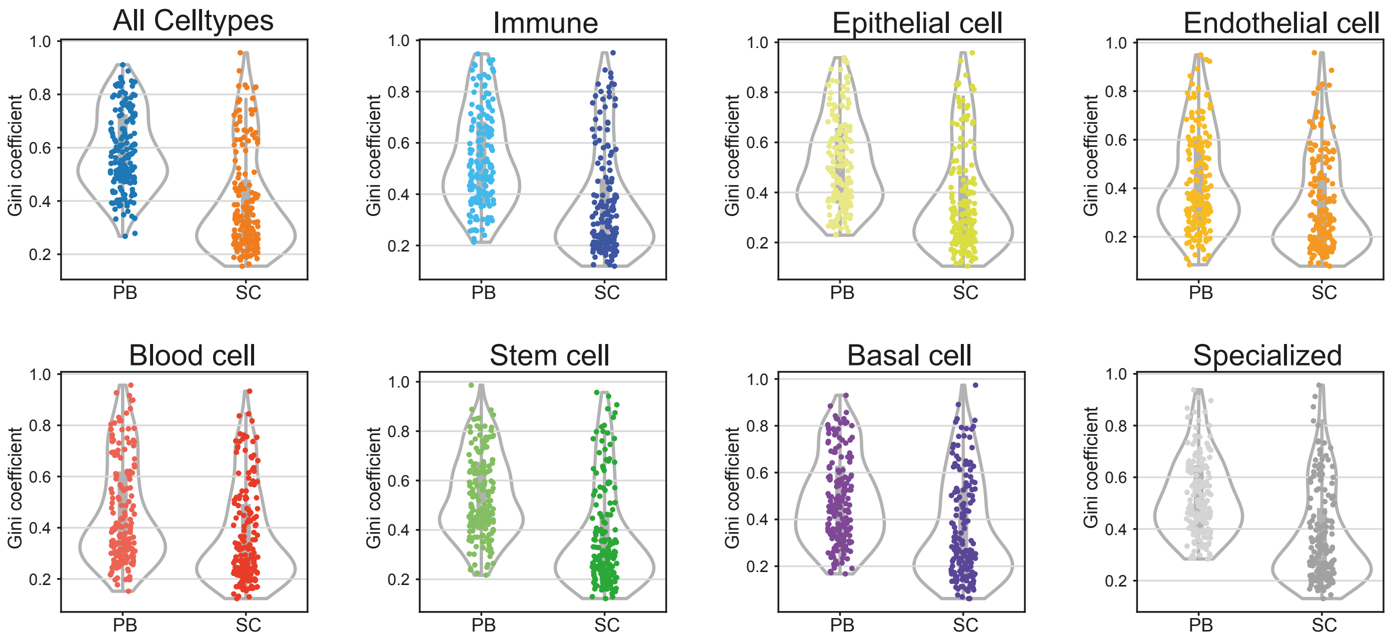


**B.**

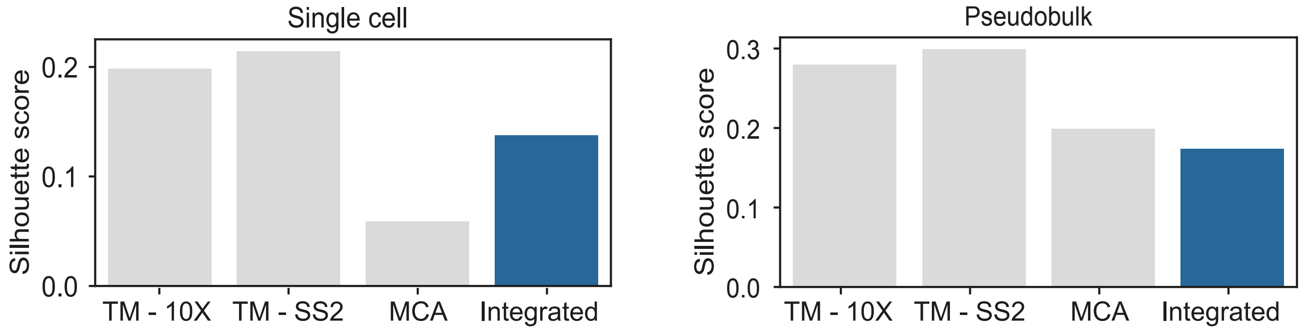


**C.** Integrated atlas (RAS correlation)

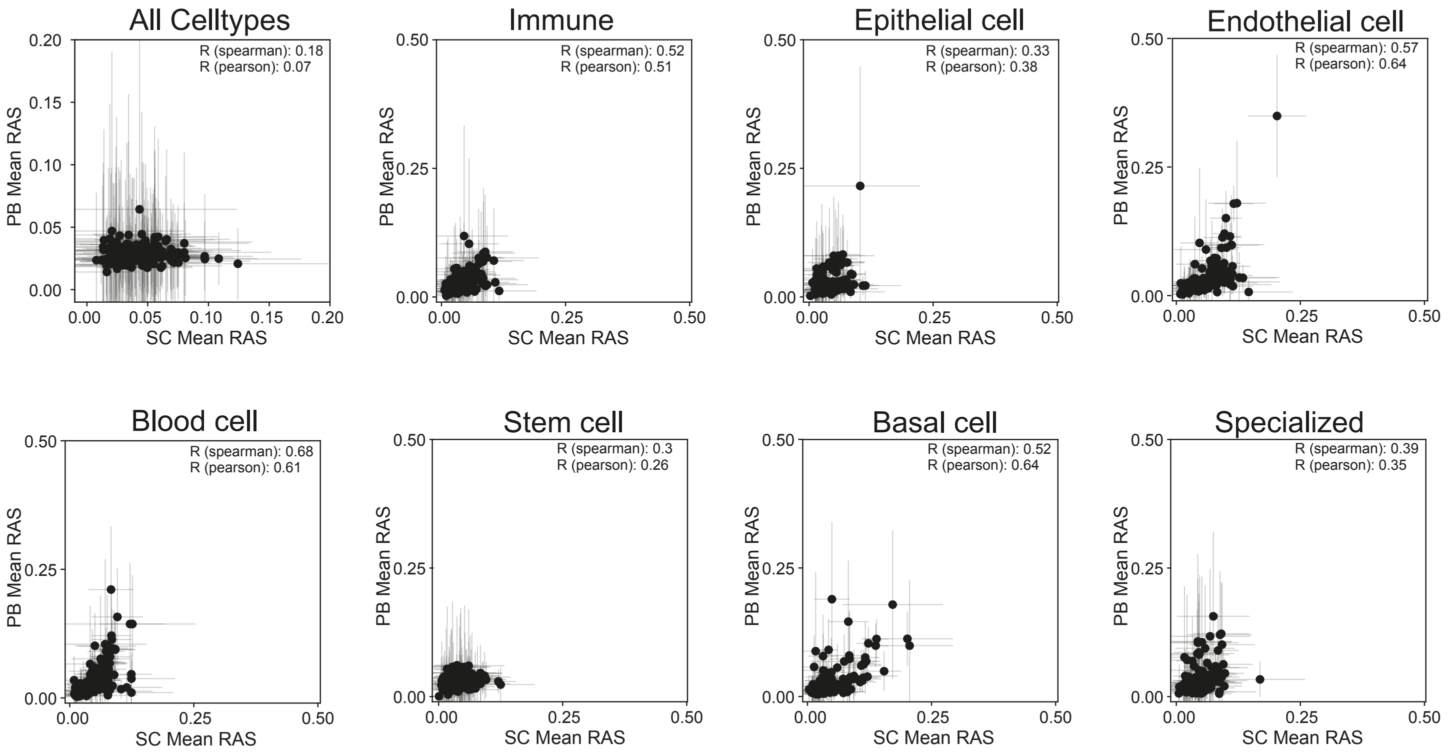

Supplementary Figure 12

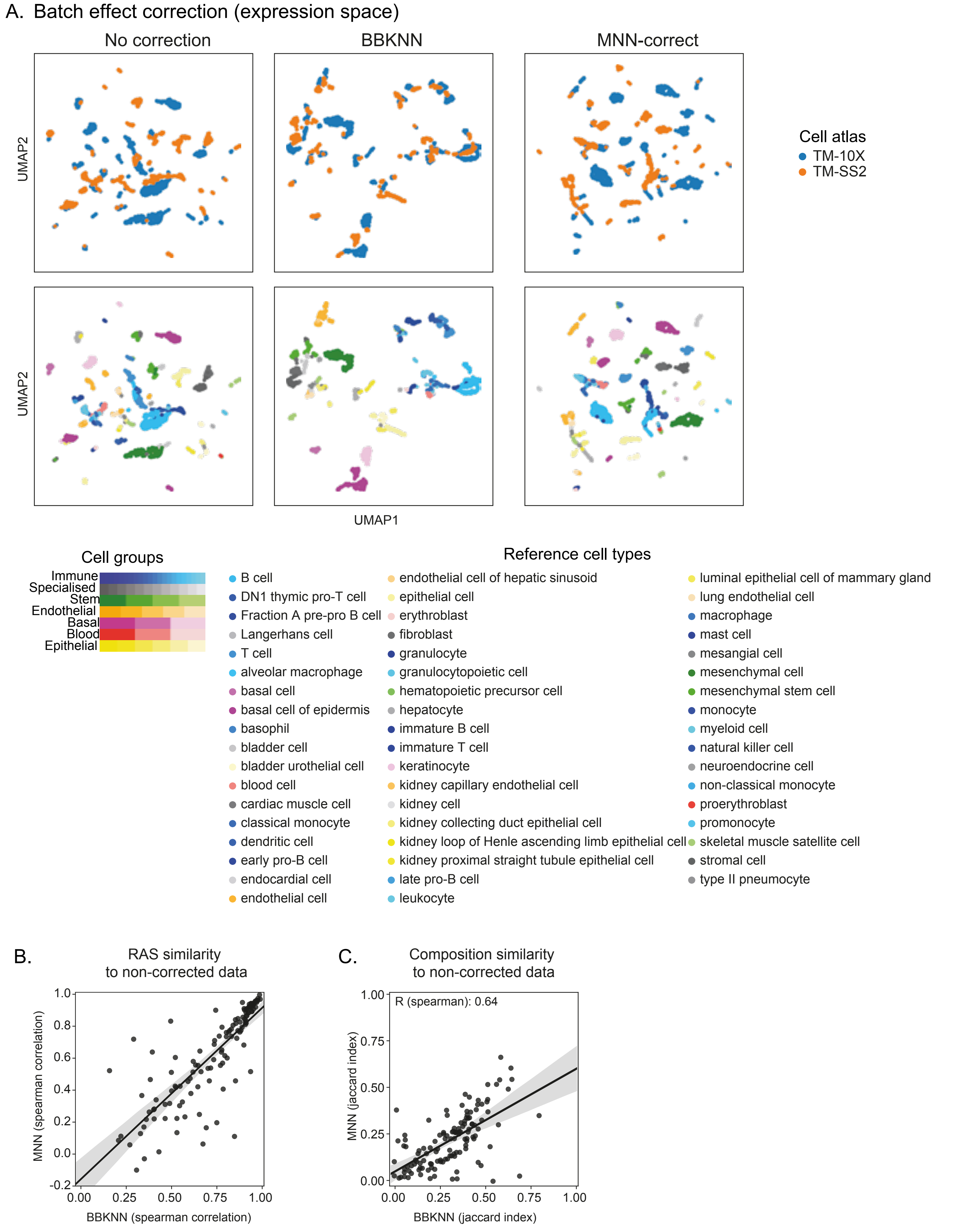

Supplementary Figure 13

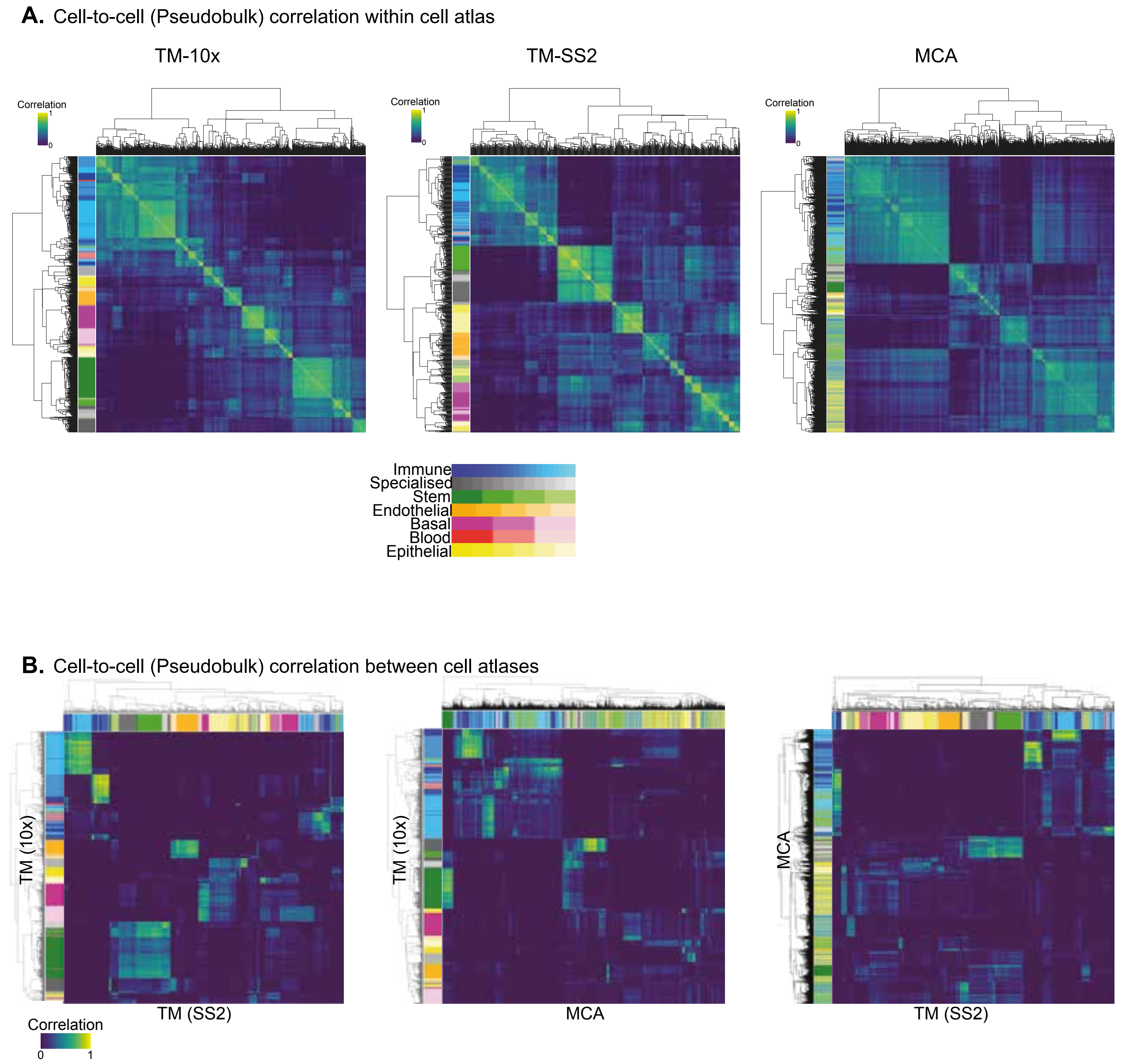

Supplementary Figure 14

**A.** Gene Ontology

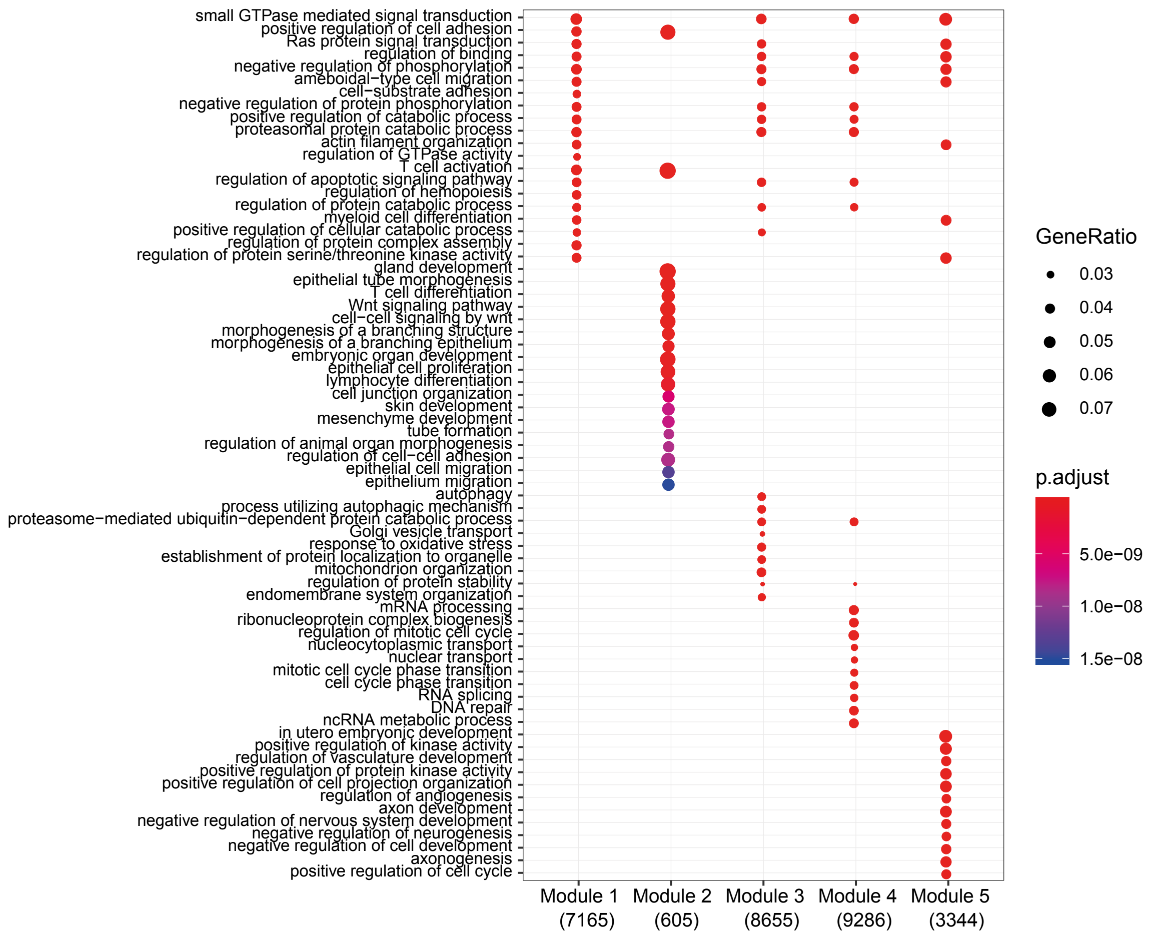


**B.** Pathway analysis (Reactome)

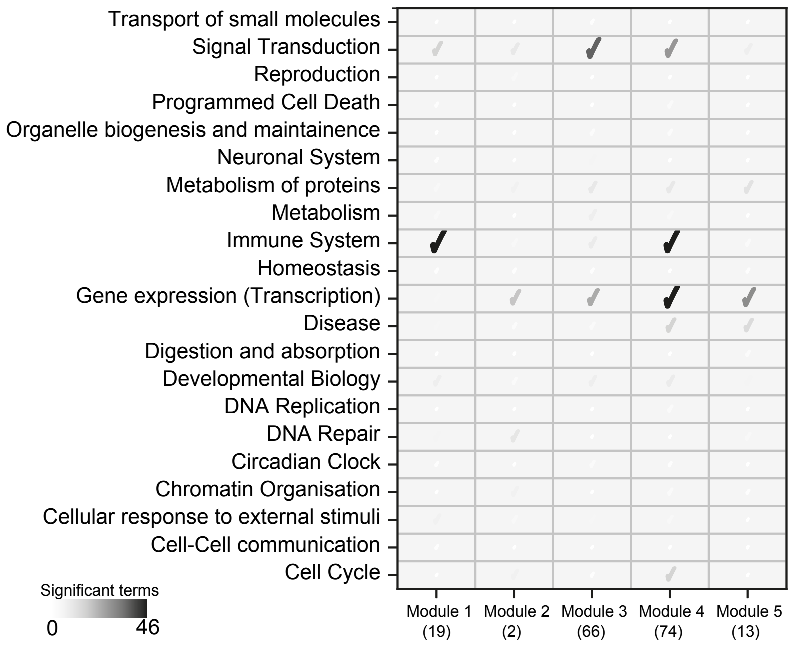

Supplementary Figure 15

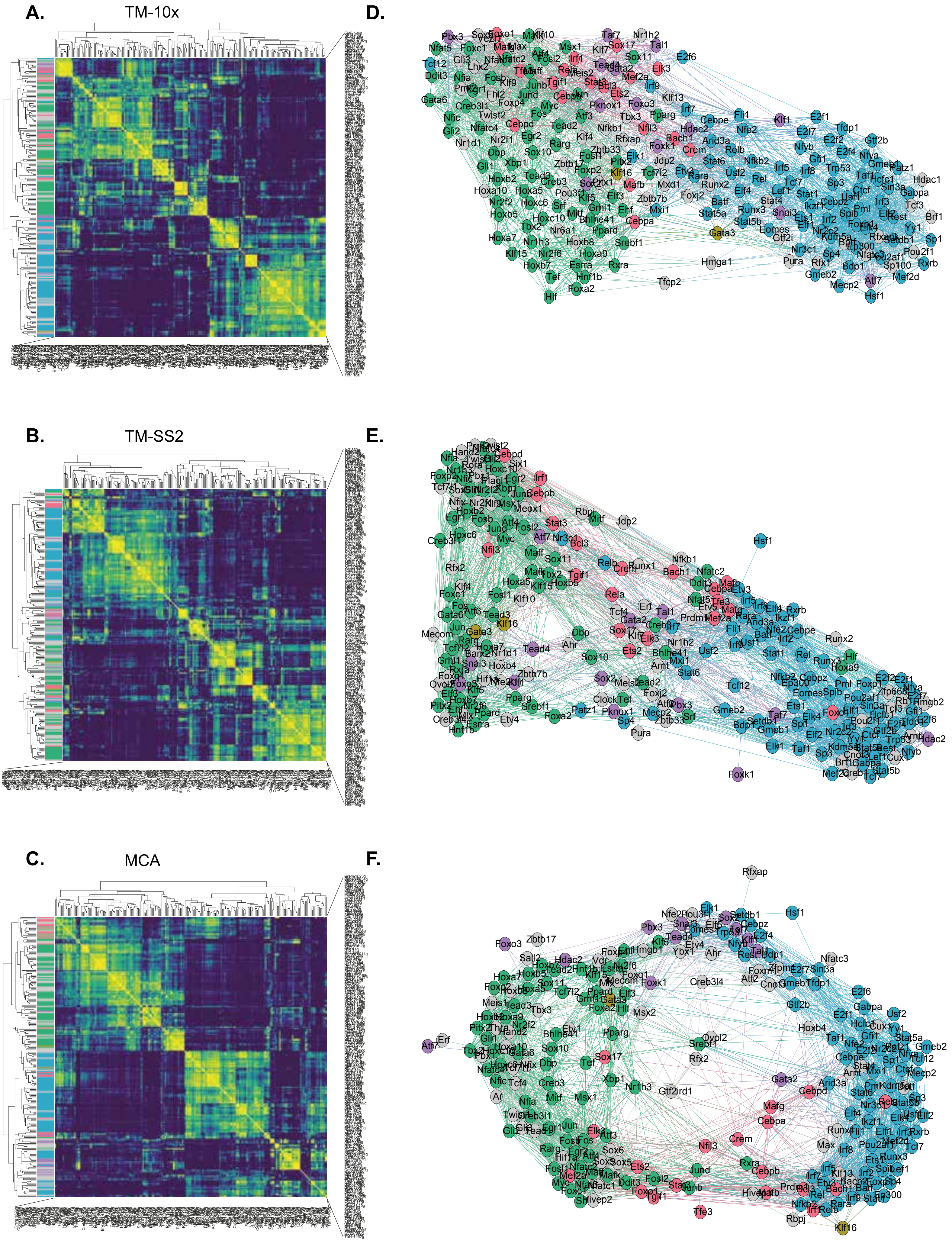

Supplementary Figure 16

**A.** TM (10x)

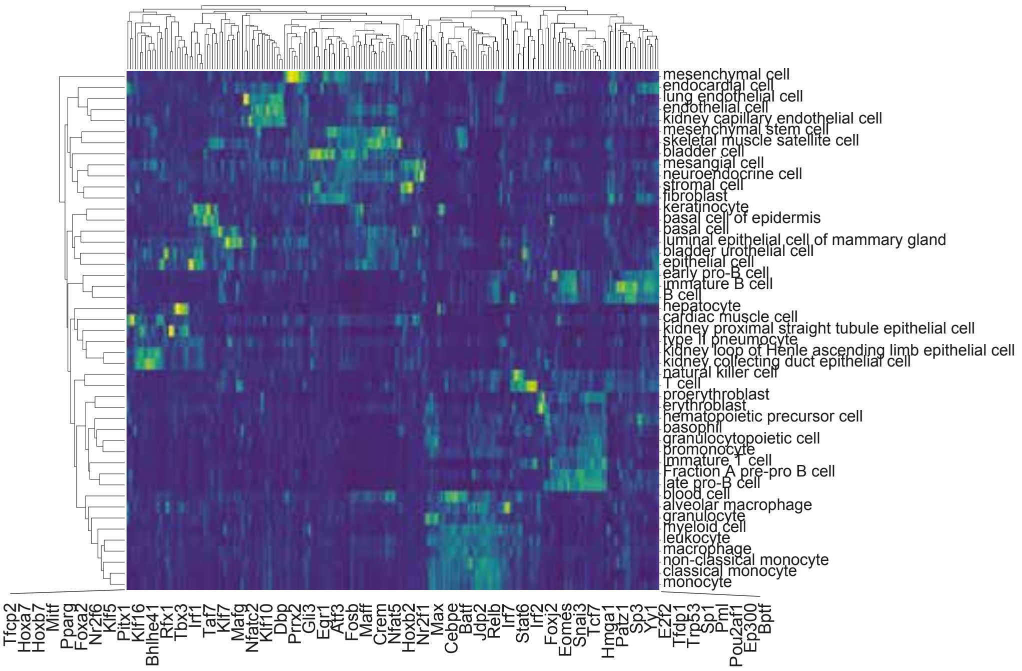


**B.** TM (SS2)

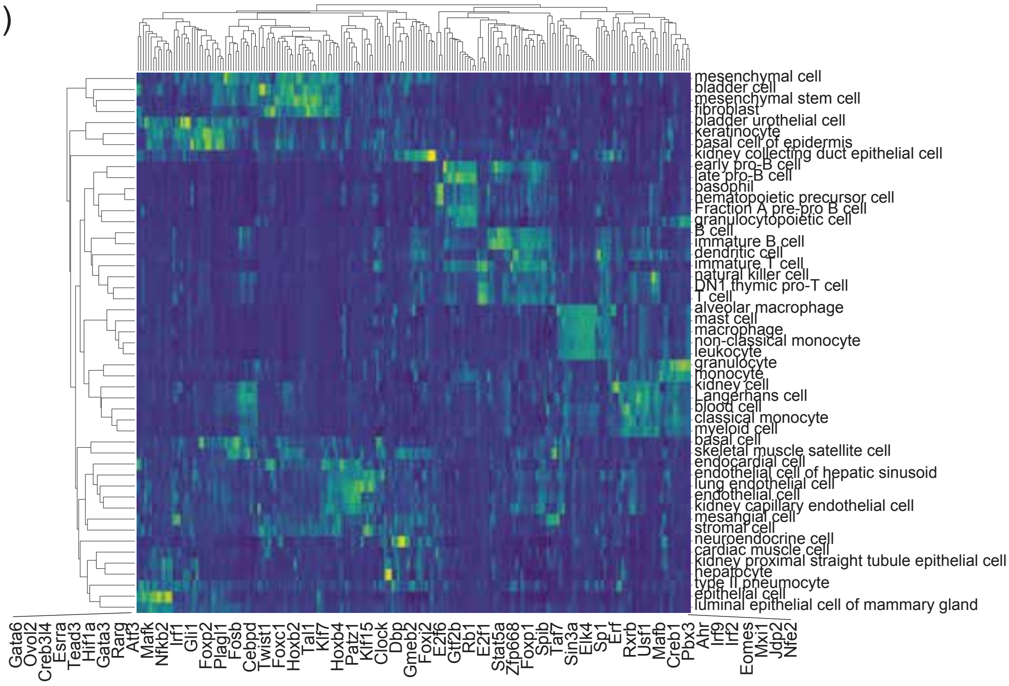


**C.** MCA

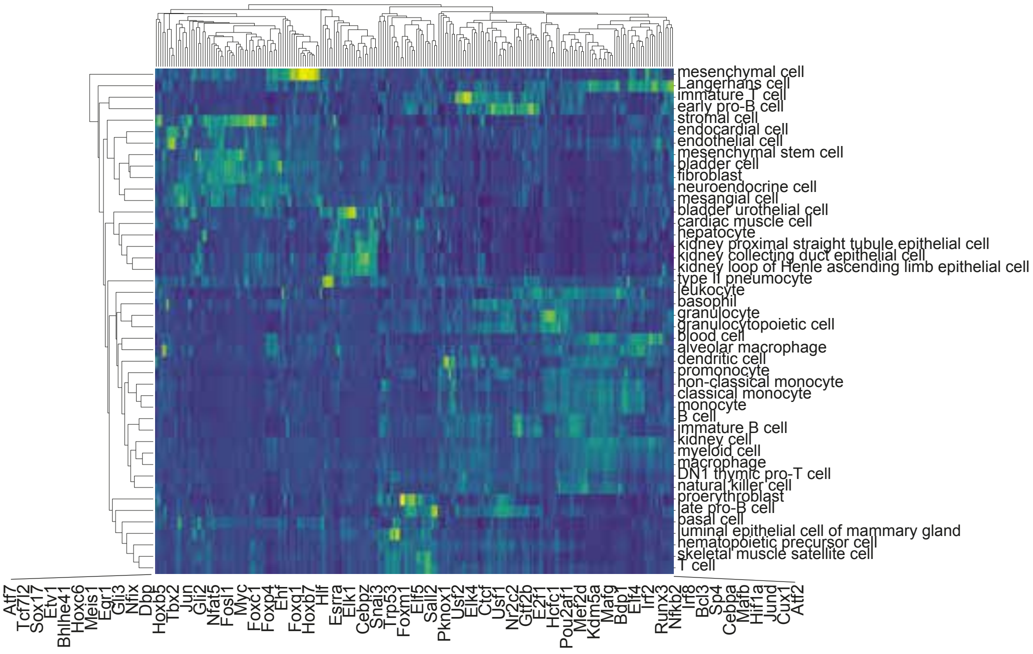

Supplementary Figure 17

A. Overrepresented motifs per module

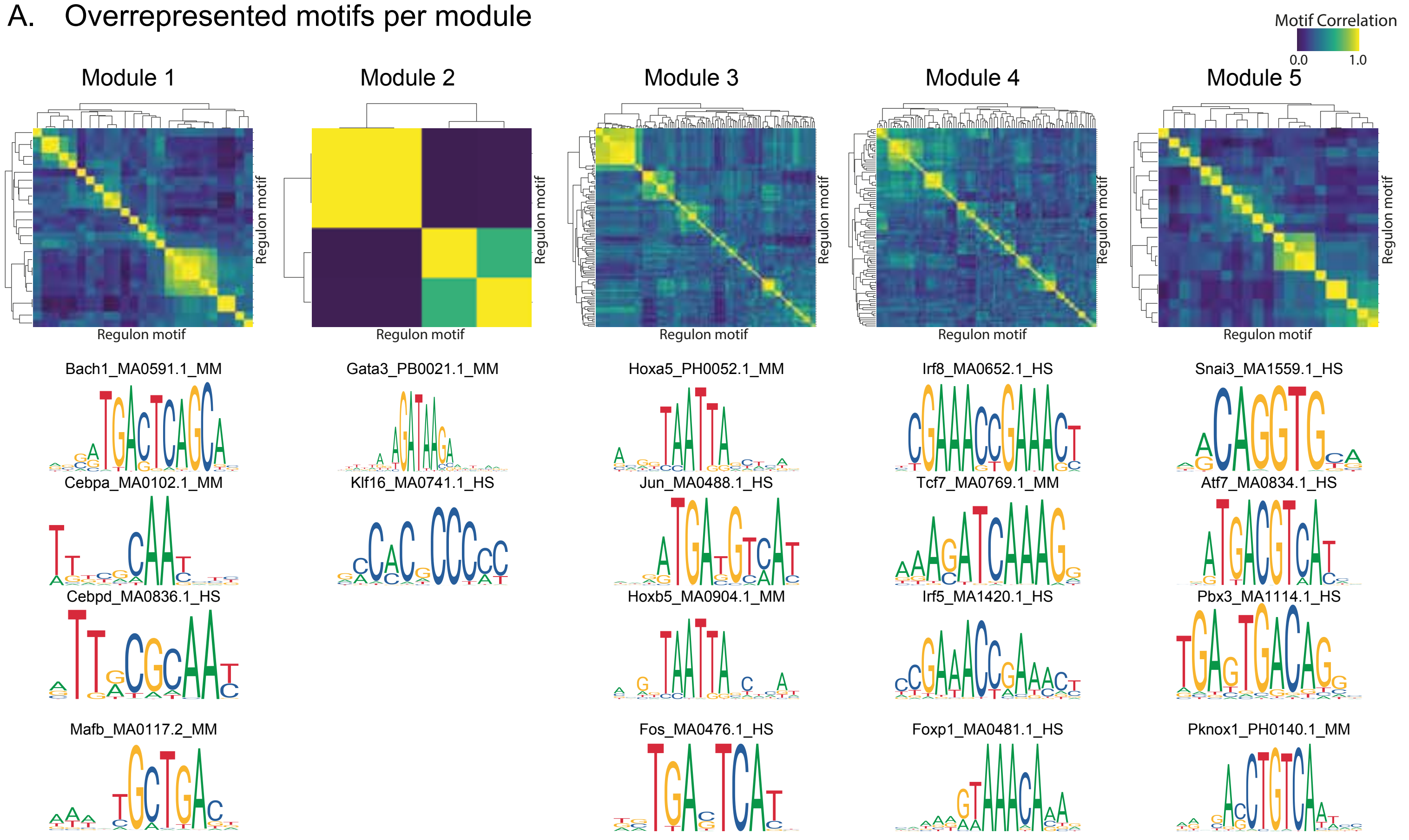


B. Protein-protein interactions (SPRING)

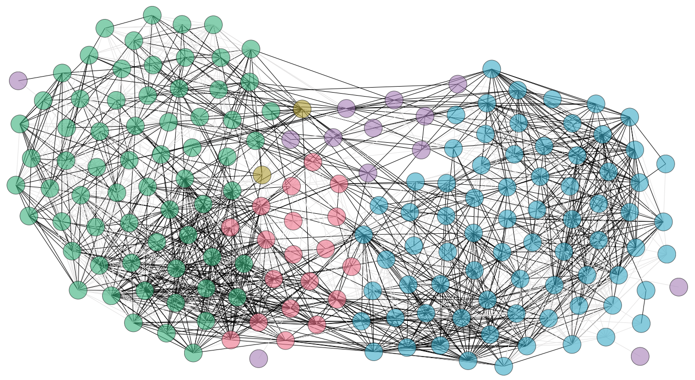

Total network edges: 4032
Edges identified in SPRING: 2290 (57%)

C.

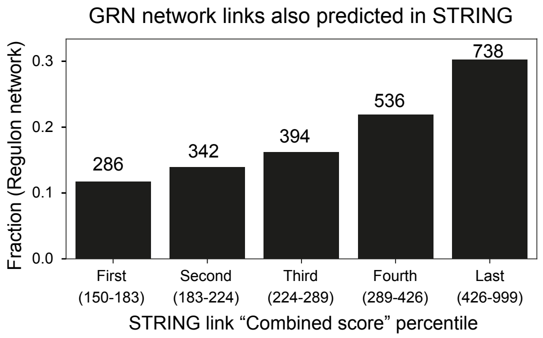


D.

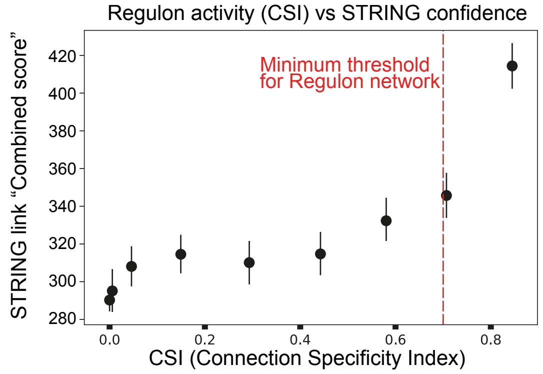


E. Essential Genes (OGEE)

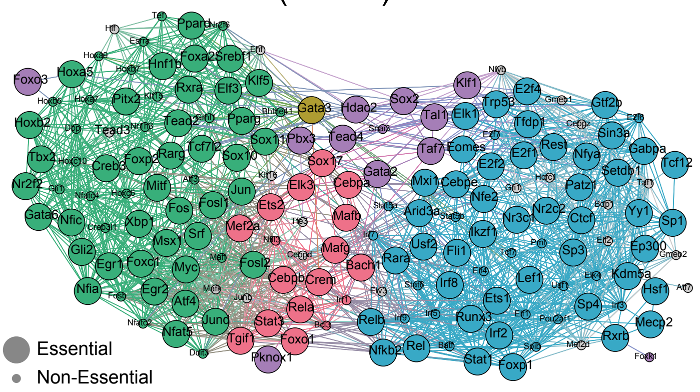


Total network nodes (regulons): 174
Network nodes in OGEE: 154 (109 essential, 45 non essential)
Network nodes not in OGEE: 20

Supplementary Figure 18

**A.** Pou2af1

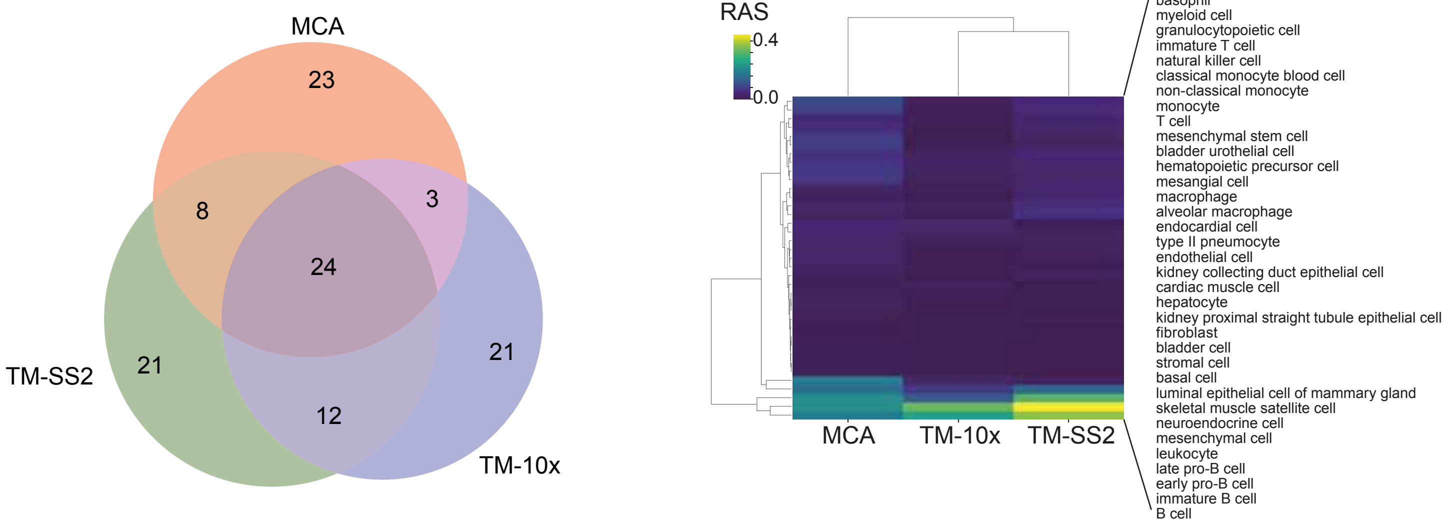


**B.** Eomes

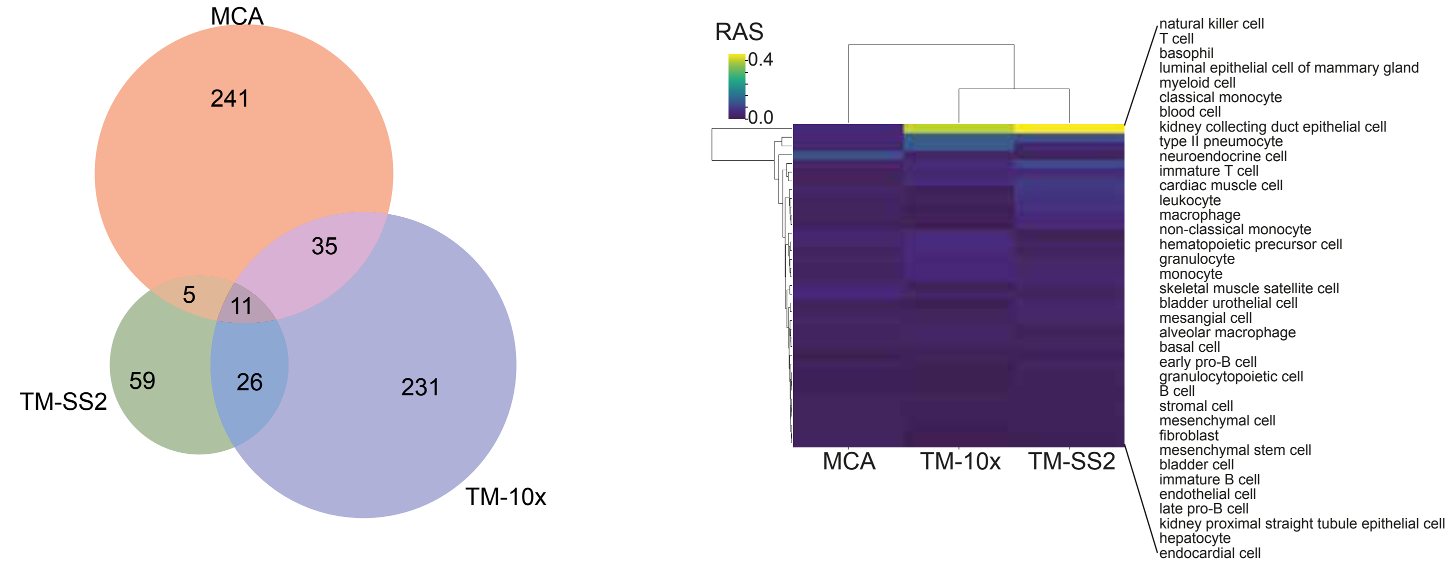


**C.** Tcf7

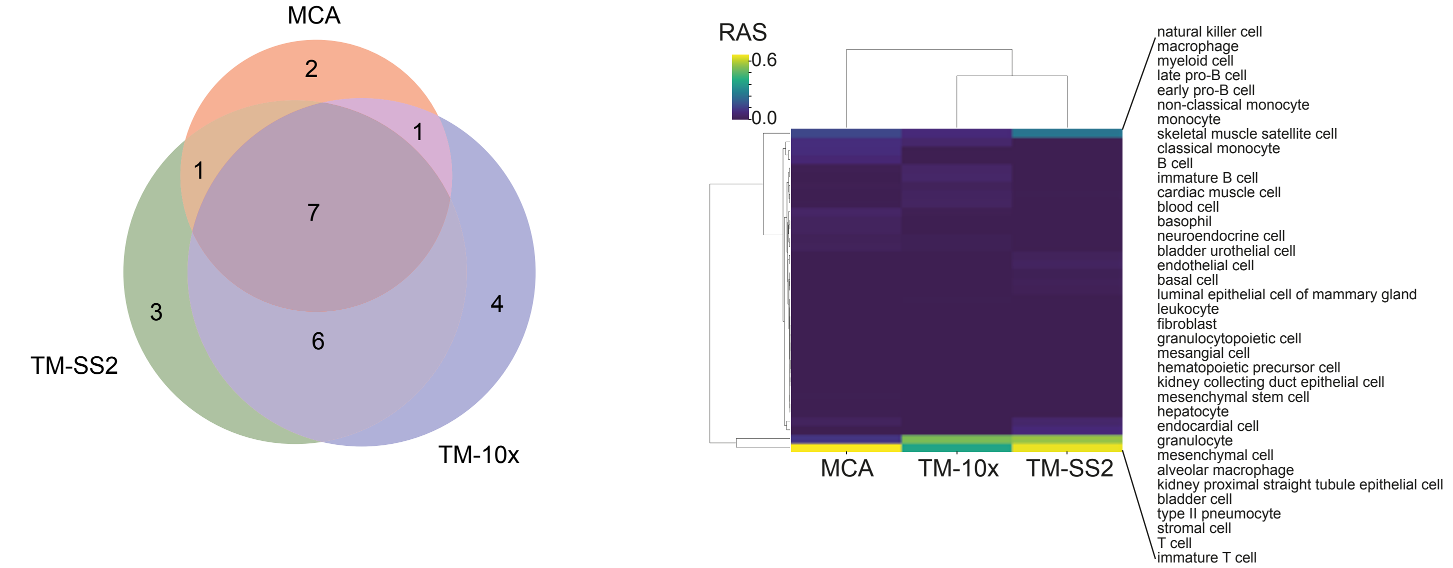

Supplementary Figure 19

**A.** Hsf1

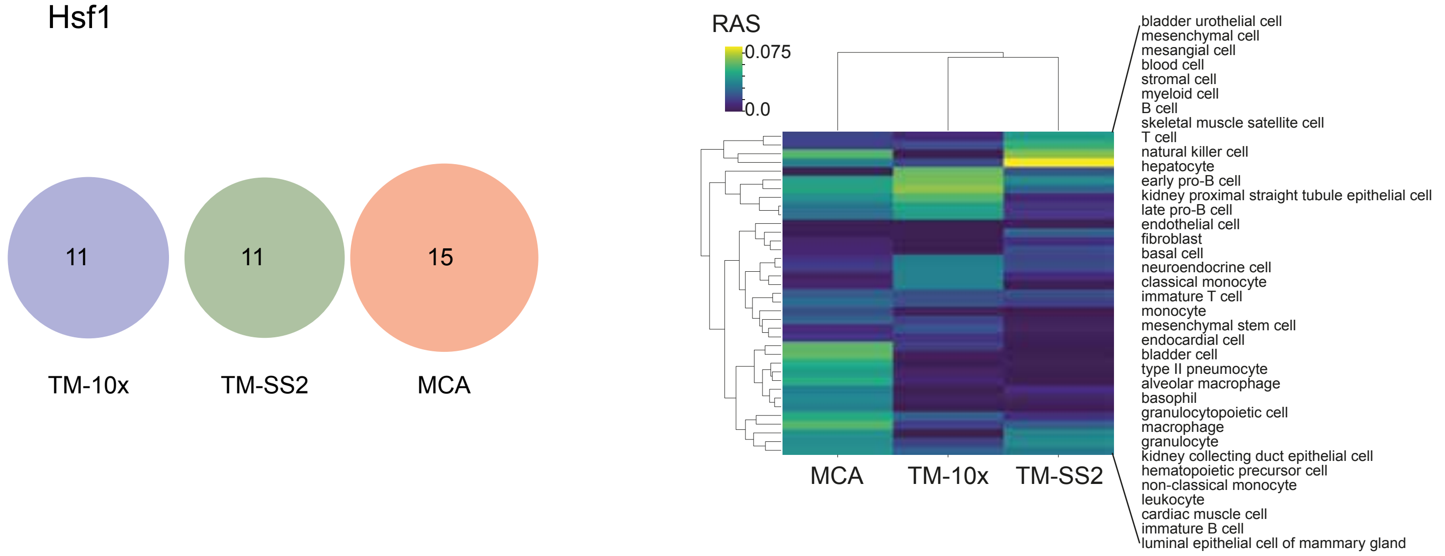


**B.** Etv3

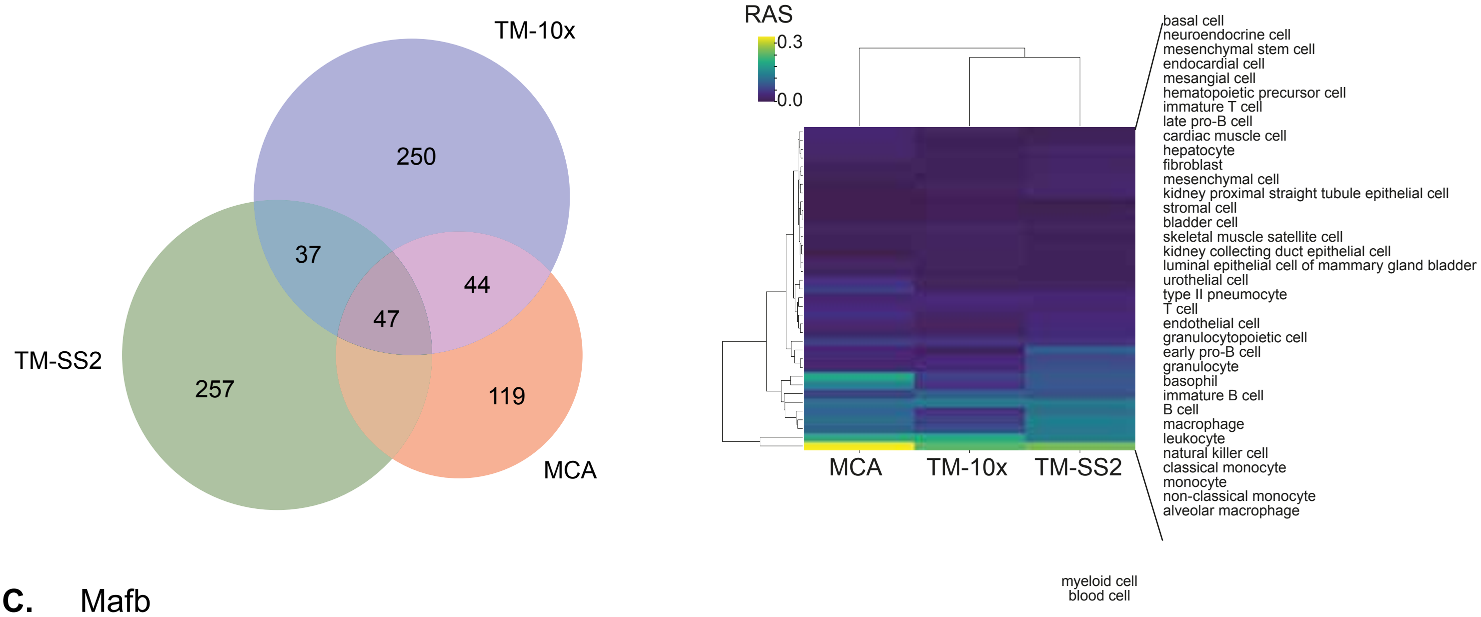


**C.** Mafb

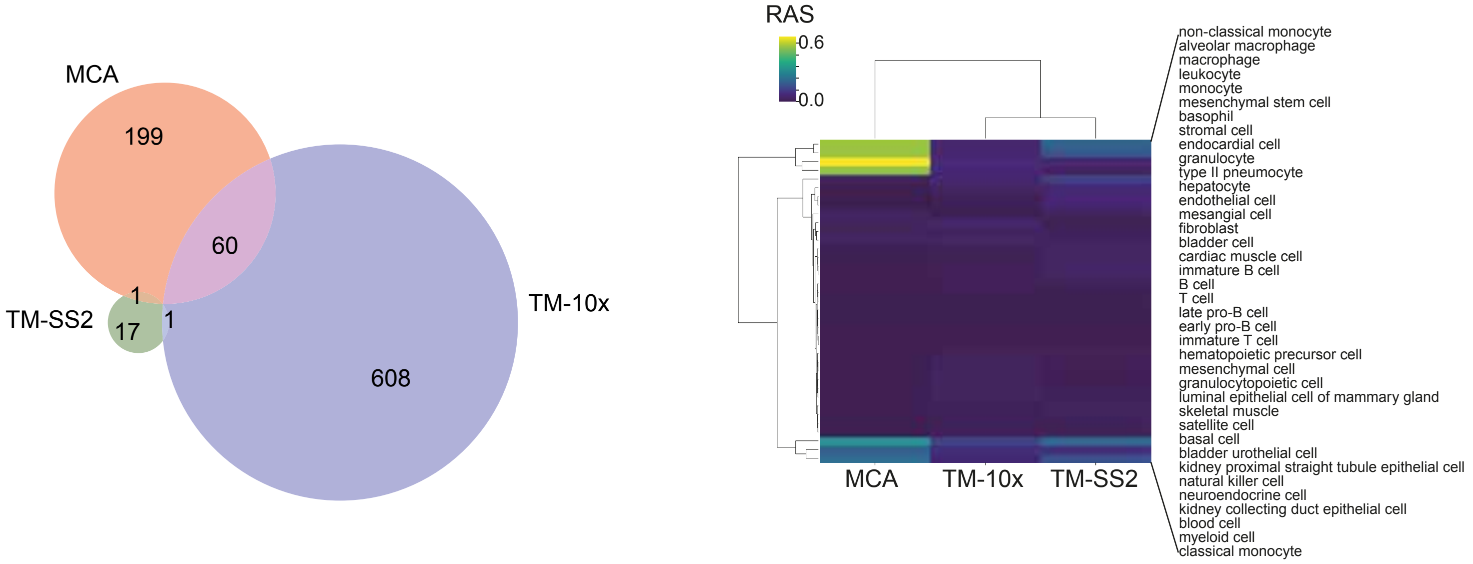

Supplementary Figure 20

**A.** Irf5

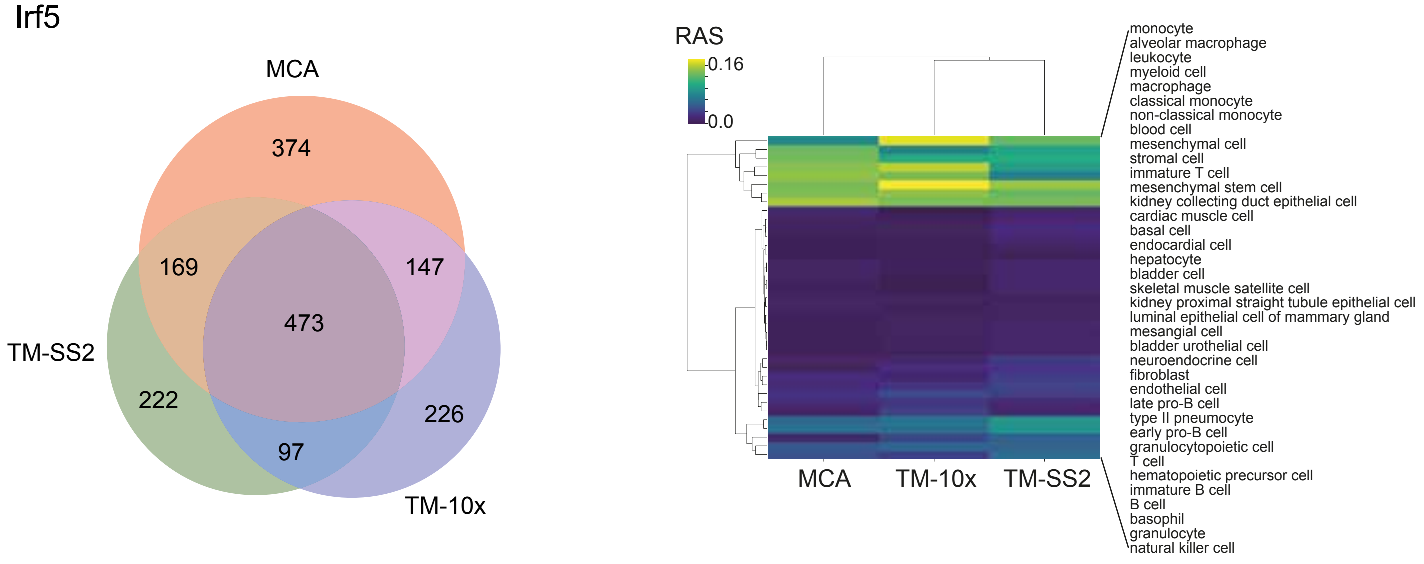


**B.** Irf9

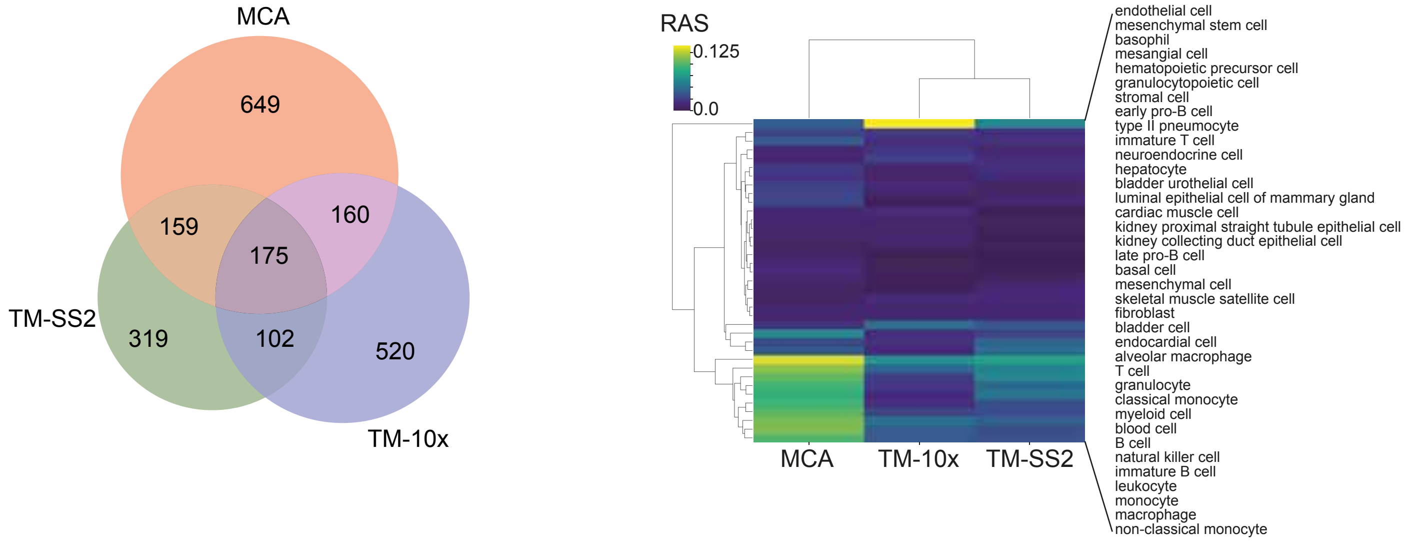


**C.** Foxp1

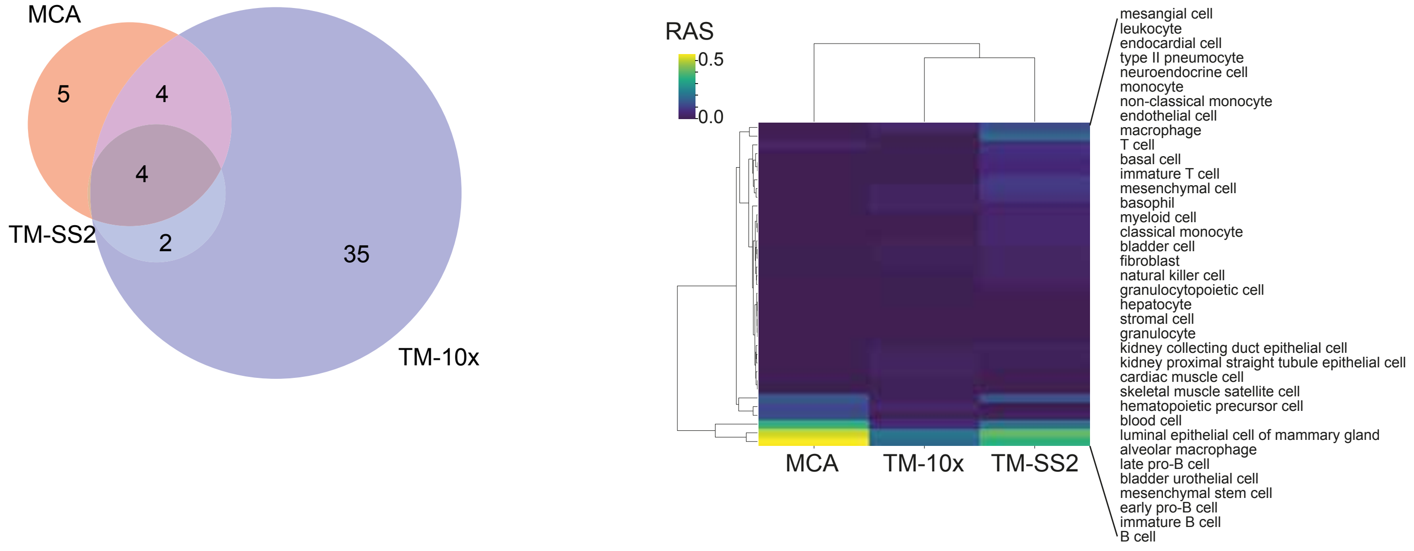

Supplementary Figure 21

A. Comparing GRN inference methods

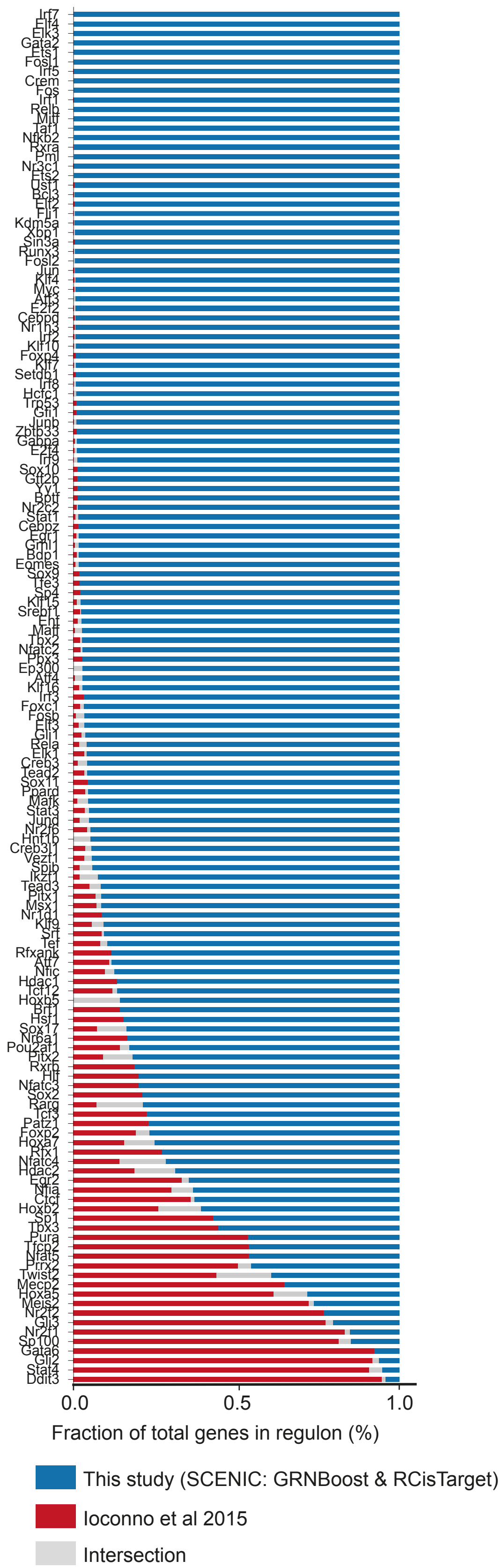


B. Regulon composition similarity

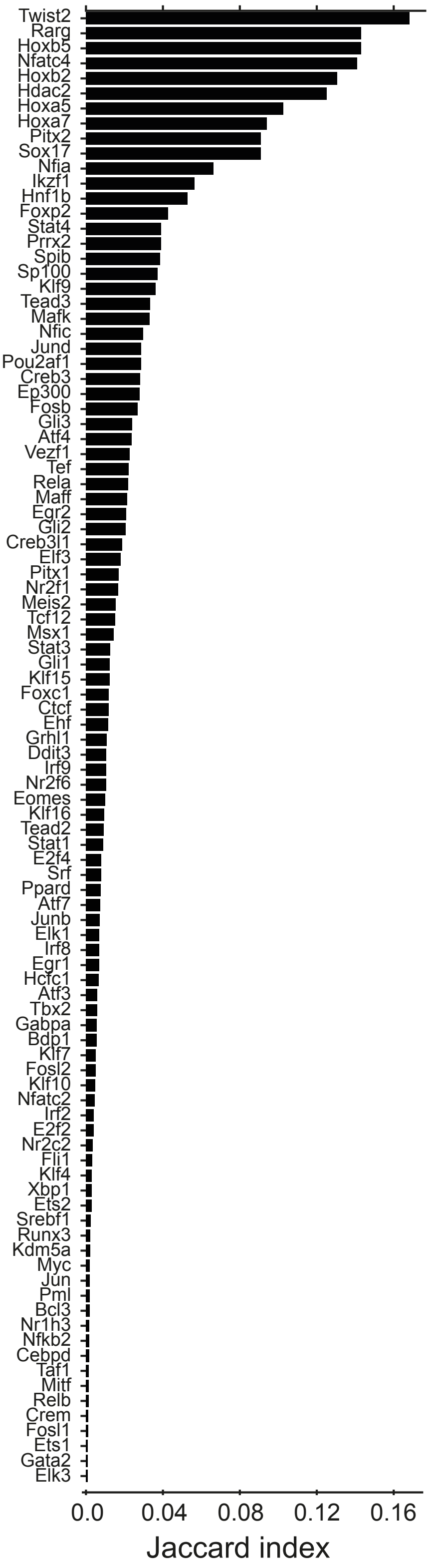

Supplementary Figure 22

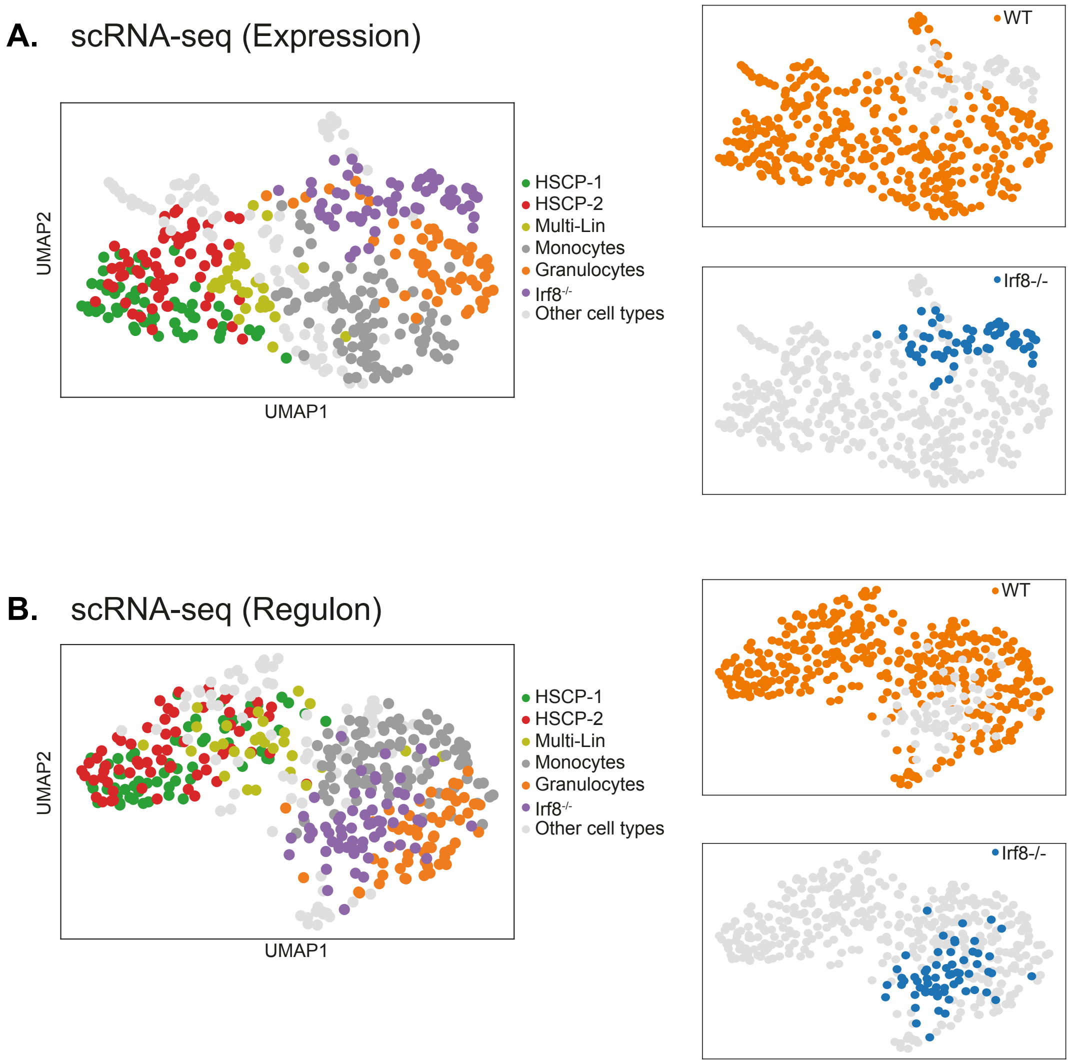


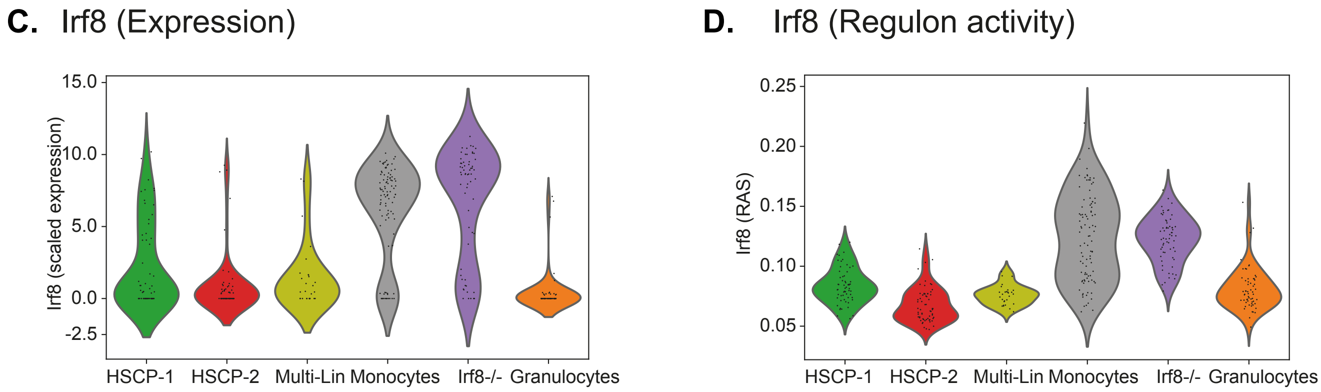


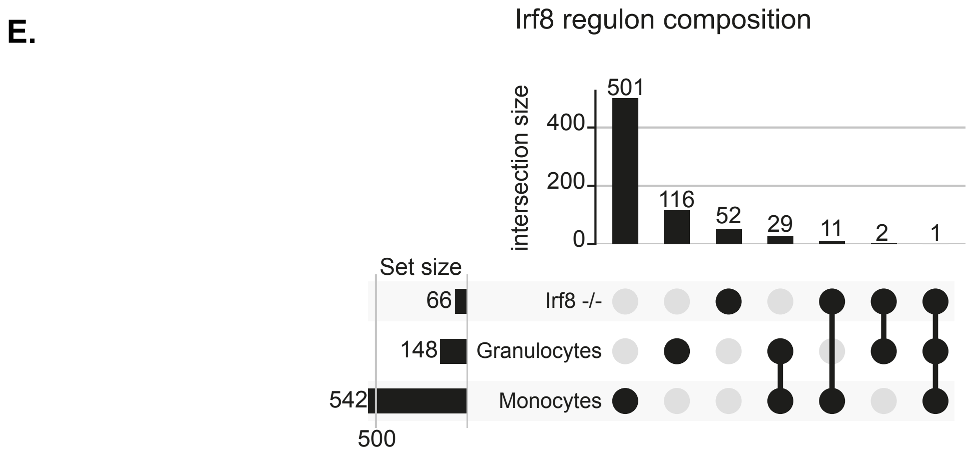
